# Single-cell transcriptomics of the immune system in ME/CFS at baseline and following symptom provocation

**DOI:** 10.1101/2022.10.13.512091

**Authors:** Faraz Ahmed, Luyen Tien Vu, Hongya Zhu, David Shing Huk Iu, Elizabeth A. Fogarty, Yeonui Kwak, Weizhong Chen, Carl J. Franconi, Paul R. Munn, Susan M. Levine, Jared Stevens, Xiangling Mao, Dikoma C. Shungu, Geoffrey E. Moore, Betsy A. Keller, Maureen R. Hanson, Jennifer K. Grenier, Andrew Grimson

**Affiliations:** Genomics Innovation Hub and TREx Facility, Institute of Biotechnology, Cornell University, Ithaca, NY 14853, USA; Department of Molecular Biology and Genetics, Cornell University, Ithaca, NY 14853, USA; Levine Clinic, Private Practice, New York, NY, USA; Workwell Foundation, Ripon, California, USA; Department of Radiology, Weill Cornell Medicine, New York, New York, USA; Department of Exercise Science and Athletic Training, Ithaca College, Ithaca, New York, USA

**Keywords:** Myalgic Encephalomyelitis/Chronic Fatigue Syndrome (ME/CFS), scRNA-seq, monocytes, platelets, Long COVID

## Abstract

ME/CFS is a serious and poorly understood disease. To understand immune dysregulation in ME/CFS, we used single-cell RNA-seq (scRNA-seq) to examine immune cells in cohorts of patients and controls. Post-exertional malaise (PEM), an exacerbation of symptoms following strenuous exercise, is a characteristic symptom of ME/CFS. Thus, to detect changes coincident with PEM, we also performed scRNA-seq on the same cohorts following exercise. At baseline, ME/CFS patients displayed dysregulation of classical monocytes suggestive of inappropriate differentiation and migration to tissue. We were able to identify both diseased and more normal monocytes within patients, and the fraction of diseased cells correlated with metrics of disease severity. Comparing the transcriptome at baseline and post-exercise challenge, we discovered patterns indicative of improper platelet activation in patients, with minimal changes elsewhere in the immune system. Taken together, these data identify immunological defects present at baseline in patients and an additional layer of dysregulation following exercise.

**Highlights:** ME/CFS is a debilitating disease with unknown causes. Here, we provide, for the first time, an extensive single cell resolution dataset detailing the gene expression programs of circulating immune cells of ME/CFS cases at baseline and after symptom provocation. We were able to detect robust dysregulation in certain immune cells from patients, with dysregulation of classical monocytes manifesting the strongest signal. Indeed, the fraction of aberrant monocytes in ME/CFS patients correlated with the degree of disease severity. Surprisingly, platelet transcriptomes were also altered in ME/CFS, and they were the only component of the immune system that showed large-scale changes following symptom provocation.

## INTRODUCTION

Myalgic Encephalomyelitis / Chronic Fatigue Syndrome (ME/CFS) is a serious human disease that lacks effective treatment options and impacts an estimated 65 million individuals worldwide (Hanson & Germain, 2020; Lim et al., 2020). Our understanding of the mechanistic basis for ME/CFS is minimal, hindering diagnosis, rational approaches to treatment options, and development of a cure. Multiple lines of evidence implicate a major role for the immune system in ME/CFS (Komaroff & Buchwald, 1998). For example, TGF-β, a cytokine that confers both pro- and anti-inflammatory signals depending on the microenvironment (Sanjabi et al., 2009), has often been reported to be upregulated in plasma of ME/CFS patients (Blundell et al., 2015; Montoya et al., 2017). Elevated levels of multiple pro-inflammatory cytokines have also been correlated with disease severity, results which suggest that ME/CFS involves dysregulation of multiple immune (and non-immune) cells. Another study observed activation of both pro- and anti-inflammatory cytokines, with different activation patterns able to differentiate early cases of ME/CFS from cases of longer duration (Hornig et al., 2015).

Indeed, immune cells of both the innate and adaptive arms are thought to be dysregulated in ME/CFS. Monocytes and natural killer (NK) cells have been reported to exhibit alterations in proportions and functional surface markers in ME/CFS (Eaton-Fitch et al., 2019; Maher et al., 2005). Neutrophils, the most abundant white blood cells in the circulation, exhibit elevated apoptosis in ME/CFS patients (Kennedy et al., 2004). T lymphocytes of the adaptive immune systems also display abnormal metabolism and cytokine production (Mandarano et al., 2020), and other studies have implicated B cell alterations in ME/CFS (Milivojevic et al., 2020; Sato et al., 2021). However, previous studies have examined different immune cell populations in isolation and often across small patient cohorts. Moreover, heterogenous populations of immune cells have typically been characterized as bulk samples, compromising the ability to distinguish changes that impact only specific cell types or subsets of cells. These limitations, together with the heterogenous nature of the disease (Committee on the Diagnostic Criteria for Myalgic Encephalomyelitis/Chronic Fatigue Syndrome et al., 2015), have likely contributed to the contradictory results between studies (Noor et al., 2021). Thus, at present, it is far from clear which components of the immune system are most relevant to ME/CFS.

A defining symptom of ME/CFS is post-exertional malaise (PEM) – an exacerbation of symptoms resulting from exertion. Thus, understanding changes that occur in one or more immune cell populations during PEM could provide useful insights to the disease, and the development of potential treatment and preventative approaches. An established method of inducing PEM in ME/CFS patients is with cardiopulmonary exercise tests (CPETs), which monitor respiratory and other parameters during a controlled exercise period with increasing intensity until exhaustion or limiting symptoms appear (Stevens et al., 2018). In addition, incorporating CPET into ME/CFS studies provides objective physiological parameters, increasing confidence in diagnosis and providing quantitative measurements of disease severity.

ME/CFS has been proposed to result from infection by an unknown virus (or other pathogen), leading to the long-lasting symptoms of the disease in susceptible individuals. This theory derives, in large part, from the many occurrences of cases of the disease in clusters (Rasa et al., 2018), together with the observation that many ME/CFS patients reported experiencing a flu-like or other infection before the onset of the disease. It is unknown whether only specific viruses can trigger ME/CFS or whether many different viruses can induce the disease, though accumulating evidence implicates the enterovirus family (O’Neal & Hanson, 2021). Notably, ME/CFS and Long COVID that arises in non-hospitalized patients share many, although not all, symptoms (Wong & Weitzer, 2021); thus, there may be important commonalities between the diseases, including a viral origin. However, the degree to which molecular signatures are shared by patients with ME/CFS and Long COVID is unknown. Therefore, knowledge gained in understanding ME/CFS may be of significance regarding Long COVID, and vice versa.

Here, we have used scRNA-seq to examine immune cells within PBMCs (peripheral blood mononuclear cells) from a large cohort of ME/CFS patients and matched controls. Our goals were to identify immune cell types with dysregulated transcriptomes in ME/CFS, and also recognize the cell types that show no substantial differences between the patient and control cohorts in order to rule out their involvement in the disease state. Finally, because PEM is a defining symptom of ME/CFS, we performed scRNA-seq both before and 24 hours after a CPET, with the goal of characterizing changes in the immune system that occur during the early stages of PEM. Amongst other changes, we observed extensive alterations in the transcriptome of a subset of monocyte cells in ME/CFS patients, with largely identical alterations present at baseline and 24 hours after the exercise challenge. Indeed, most gene dysregulation across the ME/CFS immune system was consistent between the baseline and post-exercise conditions. In contrast to this general property of the ME/CFS immune system, we observed marked differences between platelet transcriptomes from patients at baseline compared to post-CPET, suggesting that aberrant platelet activity is associated with or contributes to PEM. Taken together, this study identifies cell types with distinctive patterns of transcriptome dysregulation; these data suggest new hypotheses for understanding ME/CFS and the role of PEM in the disease.

## RESULTS

### Single-cell transcriptomics of the immune system in ME/CFS

Despite evidence that immune dysregulation is a major feature of ME/CFS, which components of the immune system are most involved in the disease is unknown. To begin to address this fundamental question, we used the 10x Genomics Chromium platform to perform single-cell RNA-seq (scRNA-seq) to profile ~5,000 PMBCs per blood sample from a cohort of 30 patients and 28 controls, matched for sex, BMI and differing in other parameters characteristic of ME/CFS (Figures 1A and 1B). Importantly, all samples were obtained prior to the Covid-19 pandemic, eliminating the possibility that any patients had Long COVID rather than ME/CFS. Because PEM is a defining symptom of ME/CFS, we profiled samples from all individuals at baseline (BL) and again 24 hours after a strenuous exercise challenge (Davenport et al., 2020; Keller et al., 2014) (post-CPET, PC; Figure 1A). This study design has the potential to define gene dysregulation in immune cells at baseline in patients as well as dysregulation associated with PEM. We used a two-step strategy to efficiently sequence each library to equivalent coverage: first, we sequenced each scRNA-seq library at low coverage (averaging 57 million reads) and used these data to determine the sequencing depth required to obtain an average depth of over 4,000 UMIs per cell, for a total average sequencing depth of 136 million reads per sample. Datasets were integrated with Seurat v4 (Hao et al., 2021) to generate a common landscape for cell type annotation and comparisons between samples. Standard scRNA-seq quality control metrics indicated robust profiling across all 30 patients and 28 control samples (Figures 1C and S1A-C; Supplemental File S1). To associate scRNA-seq clusters with cell identities, we used Seurat to systematically identify marker genes in each cluster, and we also examined expression of established PBMC marker genes, which together allowed us to identify the majority of the 28 clusters (Figures 1D and 1E).

**Figure 1.**
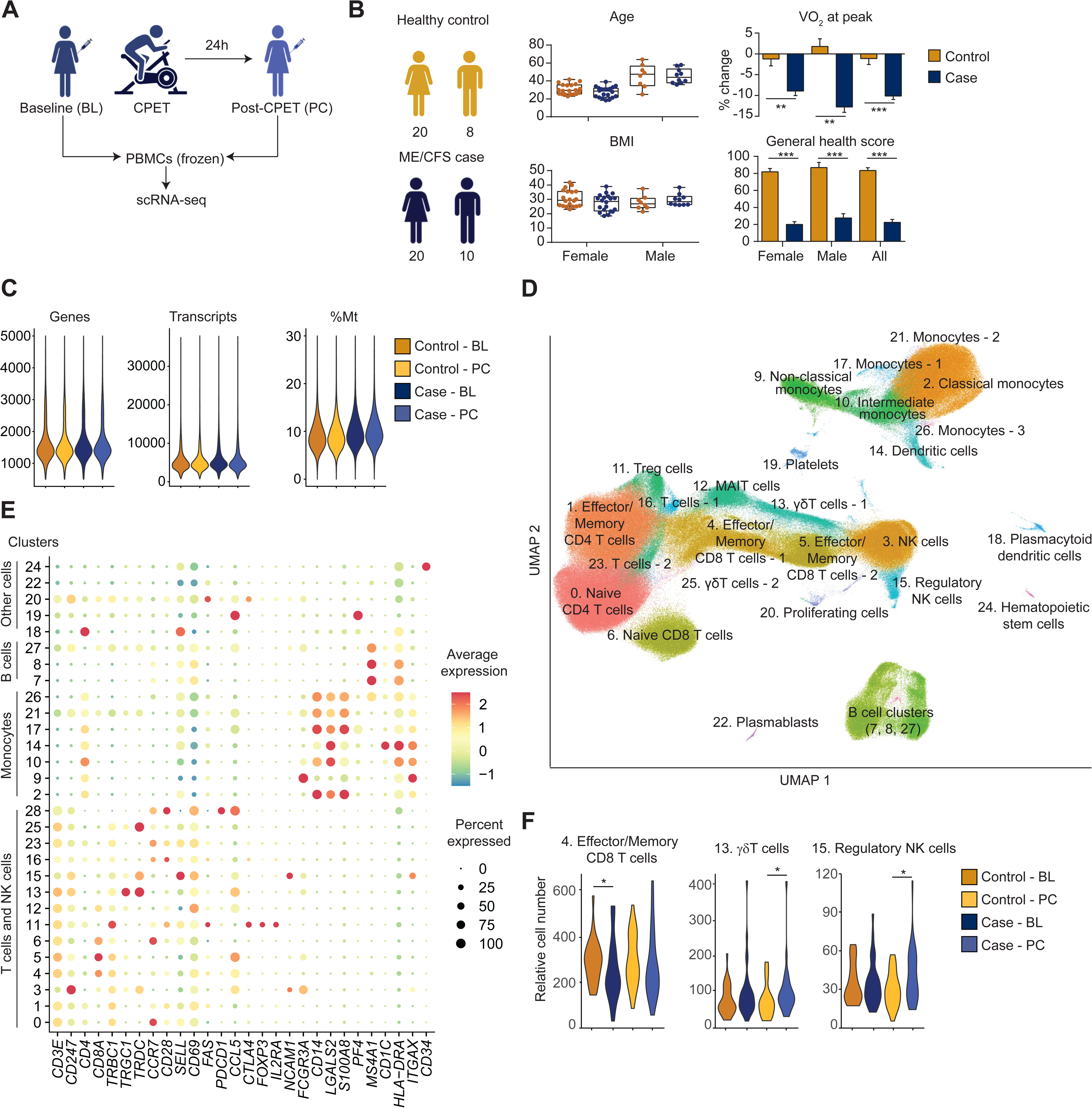
Single cell transcriptomics of the ME/CFS immune system. (A) Study design. PBMCs were collected at baseline (BL) and 24 hours post-CPET (PC) for both healthy sedentary controls and ME/CFS subjects and used for single-cell gene expression profiling (scRNA-seq). (B) Demographic and clinical parameters for patient and control cohorts. Oxygen consumption was measured during CPET. The change in maximal oxygen consumptions (VO_2_ at peak) between the VO_2_ peaks at BL and at PC is indicated. Using the SF-36v2^®^ Health Survey, general health score was self-evaluated with 100 as perfect health and 0 as worst health. Graphs represent mean ± SEM. (C) Quality control metrics, showing genes and transcripts per cell (left and middle panels, respectively) and percent mitochondrial (Mt) reads per cell (right panel), compared between four indicated cohorts. (D) Integrated UMAP with all samples. Clusters are labeled in order of decreasing number of cells with cluster 0 being the most and 27 being the least populous. (E) Relative expression of canonical and marker genes (x-axis) across clusters (y-axis), dots indicate average expression and percentage of cells with detected expression (color and size, respectively). (F) Cell types with significant differences in relative cell numbers between cohorts. *p < 0.05.

Previous studies have suggested that immune cell composition is altered in ME/CFS patients (Brenu et al., 2014; Eaton-Fitch et al., 2019; Kitami et al., 2020). Importantly, no clusters were evident that were specific to the patient cohort, nor were any largely underrepresented in this cohort (Figures S1C and S1D). We compared proportions of each cell type (cluster) between patients and controls, analyzing the baseline and post-CPET samples separately. As expected (Patel & Yona, 2019),we observed inter-individual heterogeneity in proportions of different immune cells, with these individual profiles highly consistent when compared at the two different timepoints (Supplemental File S2). However, there were only limited differences in proportions when we compared the patient and control cohorts (Figures 1F and S1D). Proportions of regulatory NK cells (cluster 15) were somewhat elevated in patients (1.1- and 1.3-fold at baseline and post-CPET, respectively; P=0.23 and 0.02, after multiple comparison correction), as were γδ T cells (1.3-fold at baseline and post-CPET; P=0.05 and 0.04, respectively). No other proportions of immune cells were significantly altered, except for a marginal decrease in effector/memory CD8+ T cells in patients (0.9-fold, at both timepoints; P=0.04 and 0.09). We conclude that cell types present in PBMCs, as profiled by scRNAseq, show very little difference in their relative frequencies in ME/CFS patients, and that any differences that do exist are minor compared to normal inter-individual variations.

### Dysregulation within the ME/CFS immune system

To begin to examine transcriptome dysregulation in the ME/CFS immune system, we used Seurat v4 (Hao et al., 2021) to determine the number of significantly dysregulated genes per cell type (cluster), comparing cells from patients and controls, and performing the comparisons separately for baseline and post-CPET samples. Certain cell types exhibit strong signals of transcriptome dysregulation in patients, with CD4+ T cells (naïve and effector/memory subsets; clusters 0 and 1, respectively), monocytes (clusters 2, 9 and 10) and cytotoxic NK cells (cluster 3) being the most prominent (Figure 2A). Importantly, there are also multiple cell types, including those present in high proportions, that show very little, if any, evidence of dysregulation in ME/CFS. We note that this analytical approach has more power to detect changes in gene expression for cell types present in higher proportions, and may have high false-discovery rates (Thurman et al., 2021). In addition, as a gauge with which to contrast dysregulation associated with ME/CFS, we also compared control samples between the two timepoints, anticipating that any signal detected for such a comparison would represent noise, at least for most cell types. This comparison revealed negligible signal, in terms of numbers of differentially expressed genes (Figure 2A), increasing confidence that the signal detected in patient versus control comparisons is meaningful.

**Figure 2.**
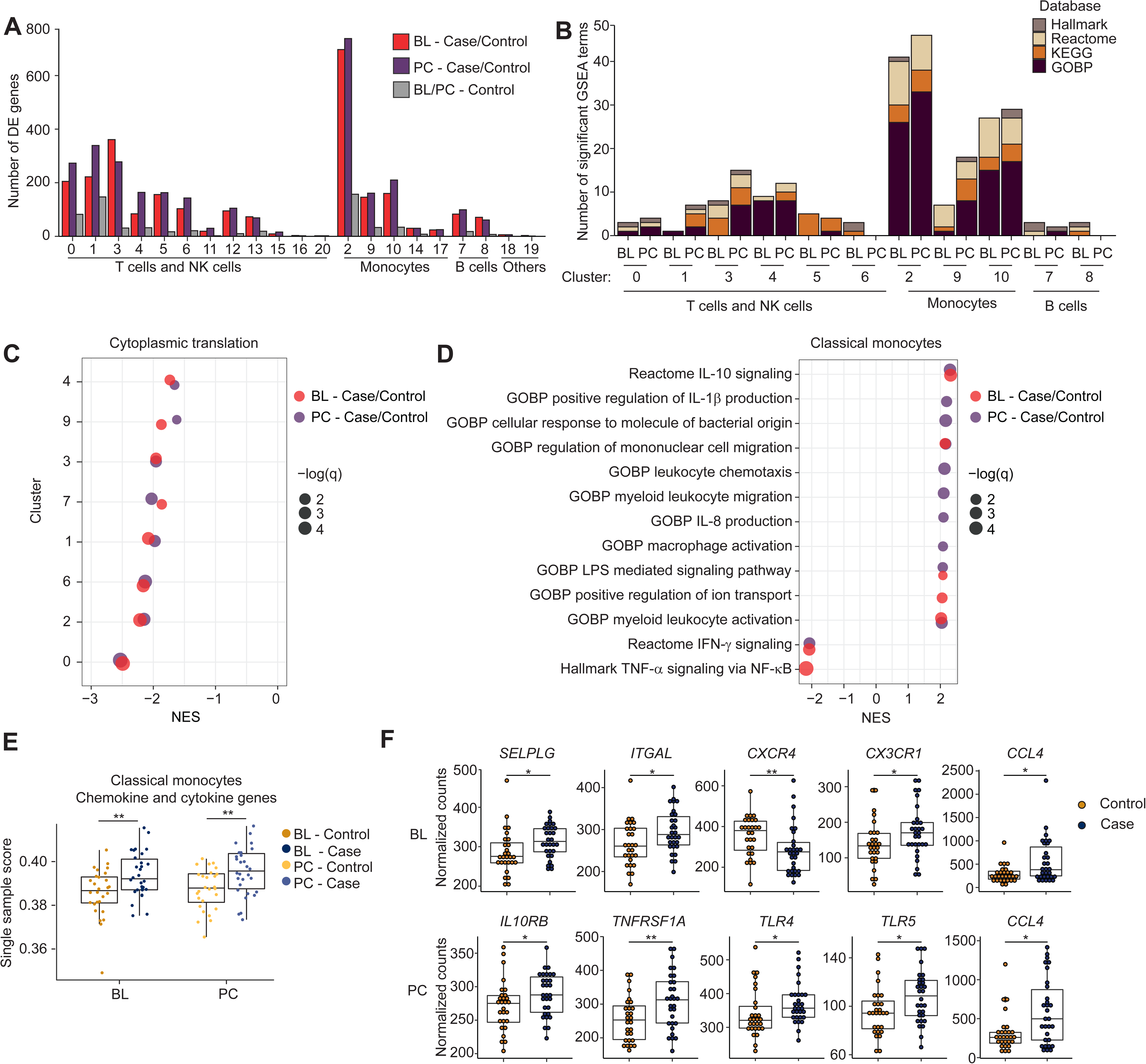
Dysregulation of immune cells in ME/CFS. (A) Counts of differentially expressed (DE) genes (y-axis) per immune cell cluster (see Figure 1 for identities), comparing case and control cells at baseline (BL) and post-CPET (PC), and compared between control cells at BL and PC. (B) Counts of the strongest significantly enriched gene sets (excluding those related to translation and with an absolute normalized enrichment score (NES) less than 2) across the largest immune cell clusters. (C) Representative GSEA gene set (GOBP cytoplasmic translation) showing lower detection of ribosomal proteins in major clusters (NES < 0) at both BL and PC in ME/CFS (red and purple, respectively). Dots are sized to denote significance (q-values); x-axis indicates NES. (D) GSEA results for classical monocytes (cluster 2) comparing patient and control cohorts at baseline and post-CPET (red and purple, respectively), focusing on gene sets related to chemokine/cytokine signaling. Dots are sized to denote significance; x-axis indicates NES. (E) Single sample scores generated using GSEA of leading-edge genes from panel D. ** p < 0.01. (F) Differential expression of genes associated with monocyte migration and differentiation at baseline and post-CPET in classical monocytes. * p < 0.05, ** p < 0.01.

Multiple frameworks exist with which to detect differential gene expression in scRNA-seq. Recent approaches have indicated that ‘pseudobulk’ methods (aggregating expression signals across all cells per cluster) can outperform the Seurat framework (Thurman et al., 2021). However, we found that this approach had limited power to identify individual genes with significant differences between case and control samples in different cell types, likely because inter-individual variation in gene expression unrelated to disease state is a large confounding factor, and the observation that the largest variation in the dataset can be attributed to sex (Figure S2A). Indeed, previous studies have identified sex-specific changes in microRNA profiles from PBMCs in ME/CFS (Cheema et al., 2020). Instead, to search for coordinated shifts in gene expression reflecting alterations in pathway activation or cell state in the ME/CFS immune system, we performed GSEA (gene set enrichment analysis) on each cluster, focusing on C1: HALLMARK, C2:REACTOME, C2:KEGG, and C5:GOBP (biological process) catalogs from the Molecular Signatures collection (Liberzon et al., 2011; Subramanian et al., 2005). This analysis (Figure 2B; Supplemental File S3) indicated two general features of the data: first, across multiple clusters, expression of ribosomal protein genes and core translational machinery is downregulated in ME/CFS relative to controls in many different cell types (Figure 2C), and second, that monocytes exhibited the strongest signals of dysregulation, especially in cluster 2 (classical monocytes) but also in clusters 9 and 10, which represent non-classical and intermediate monocytes, respectively (Figure 2B, Figure S2B; Supplemental File S4). In particular, gene sets associated with chemokine signaling, migration and activation are expressed at elevated levels in classical monocytes from patients (Figure 2D). In addition, we also observed suppression of genes associated with interferon gamma (IFN-γ) signaling in classical monocytes from ME/CFS patients. These results suggest that the transcriptomes of classical monocytes from ME/CFS patients are biased towards a profile that promotes migration of monocytes to tissue and increased progression towards a macrophage fate. However, we also observed activation of genes associated with IL-10 signaling, which is an anti-inflammatory cytokine. Thus, monocytes in ME/CFS patients, and perhaps also the macrophages derived from such monocytes, might undergo a combination of conflicting signaling inputs.

To explore signals that might be differentially impacting monocytes in patients, we examined the identities and expression levels of the genes driving the enrichments we observed. Using the genes most responsible for enriched gene sets related to chemokine and cytokine signaling (leading edge genes; Supplemental File S5), we calculated a composite score for classical monocytes from each sample (Figure 2E; other monocyte clusters are shown in Figure S2C). At the level of individual samples, we found classical monocytes from ME/CFS cases to have higher scores for these genes compared to sedentary healthy controls, and that these scores are consistent between baseline and post-CPET timepoints for each individual (Figure S2D). We also looked at differential expression of multiple notable genes involved in monocyte response, recruitment and differentiation at baseline and post-CPET. Specifically, we observed upregulation of *CCL4*, *CX3CR1*, *SELPLG* and *ITGAL* in patients, while *CXCR4* expression was suppressed at baseline (Figure 2F). CCL4 is a chemoattractant for monocyte recruitment to inflamed and adipose tissue (Sindhu et al., 2019). CX3CR1, the sole chemokine receptor for CX3CL1, is a marker for differentiation of classical monocytes to non-classical counterparts and tissue repair and regeneration (Feng et al., 2015; Getzin et al., 2018). SELPLG and ITGAL play critical roles in the tissue recruitment of leukocytes (Muller, 2013). CXCR4 is a receptor for CXCL12 and regulates monocyte-macrophage differentiation (Sánchez-Martín et al., 2011). *CCL4* was also upregulated regardless of exercise challenge. We also detected upregulation of both inflammatory (*TNFRSF1A, TLR4, TLR5*) and anti-inflammatory (*IL10RB*) receptor genes post-CPET. Taken together, these results suggest that monocytes in ME/CFS are aberrantly pro-migratory and inflammatory at baseline and that ME/CFS is characterized by a persistent state of monocyte activation.

### Transcriptome profiling of purified classical monocytes from ME/CFS patients and controls

To generate a more comprehensive profile of dysregulation of classical monocytes in ME/CFS and to validate our scRNA-seq results, we turned to bulk RNA-seq of purified monocytes. Starting with PBMCs from four female patients and four female controls isolated post-CPET, with all individuals distinct from those profiled by scRNA-seq, we isolated near-homogenous populations of classical monocytes (CD14+CD16-). Flow cytometry confirmed that the isolation strategy was effective (Figure 3A). Following RNA isolation, we generated RNA-seq libraries sequenced to a minimum depth of 20 million reads per sample. Principal component analysis demonstrated clear separation between patient and control samples, although the patient samples were more dispersed than controls (Figure 3B). We compared expression profiles of classical monocytes from patients and controls (Figure S3A) and identified enrichment of similar GSEA terms and leading-edge genes between the bulk RNA-seq (Figure 3C and Supplemental File S6) and the scRNA-seq (Figure 2D and Supplemental File S4), such as regulation of cell migration by cytokines and chemokines and IL-10 signaling. These observations confirmed a pattern of monocyte activation and migration in ME/CFS in comprehensive expression profiles of an independent cohort of samples. We also examined expression levels for a large set of genes associated with monocyte migration and differentiation in the bulk RNA-seq (Figure 3D and S3B). Perhaps due to heterogeneity in the ME/CFS cases (Figure 3B), we found consistent changes only in *CCL4* upregulation and *CXCR4* downregulation, when examining both the bulk RNA-seq for classical monocytes and the corresponding cluster from the scRNA-seq (Figures 2F and 3D). However, using the RNA-seq data alone, we identified additional changes in expression, including most prominently upregulation of *CSF2* (granulocyte-monocyte colony stimulating factor; an important monocyte differentiation cytokine) and other chemokines important for monocyte recruitment (*CCL20, CCL5,* and *CCL3*) (Zhao et al., 2020) in ME/CFS cases. These analyses support the hypothesis that monocytes in ME/CFS patients are aberrantly primed to migrate to tissues.

**Figure 3.**
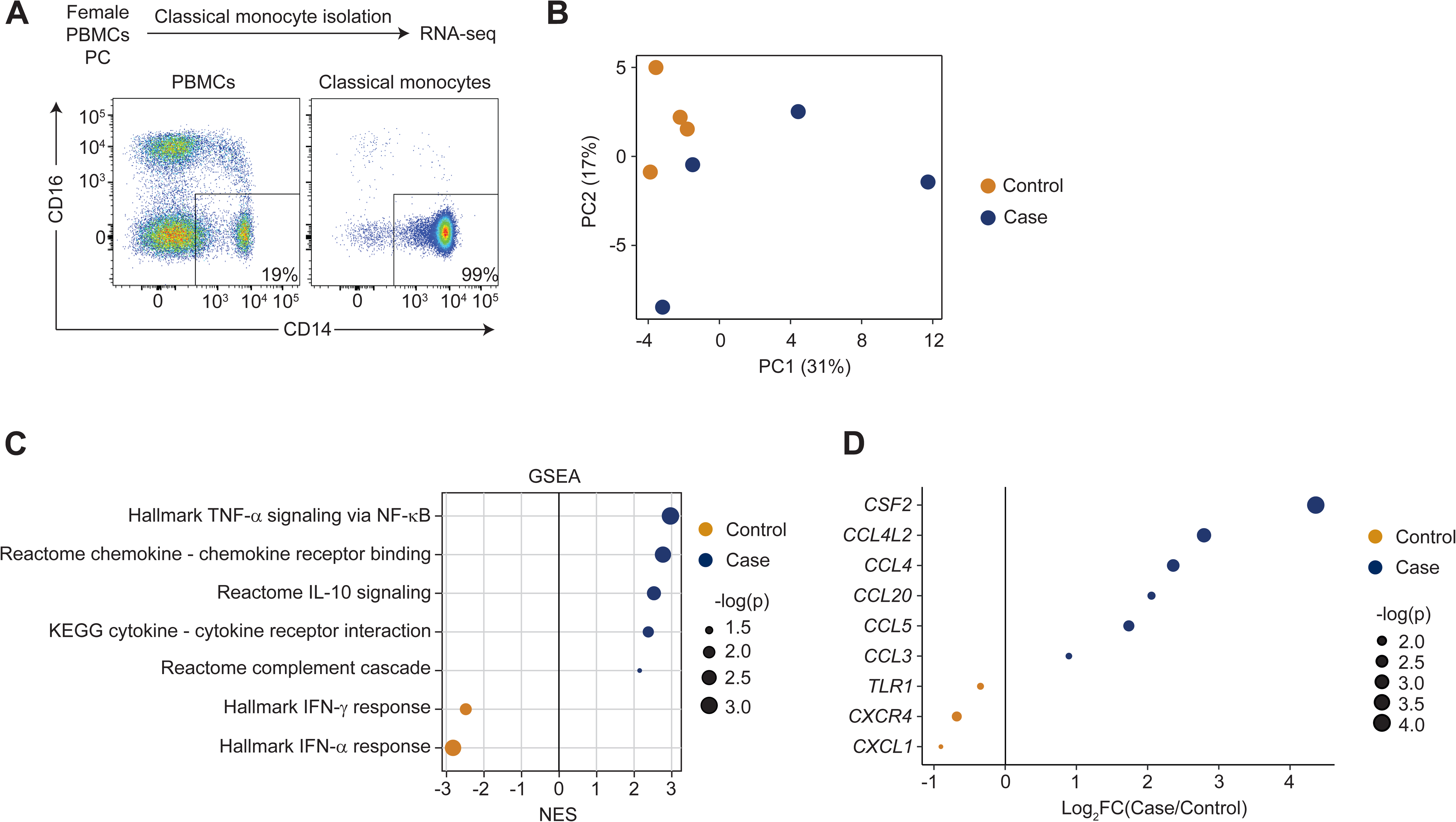
Complete transcriptomics of classical monocytes. (A) Purification of classical monocytes from PBMCs. PBMC samples from female cases and controls collected post-CPET was utilized for classical monocyte isolation and bulk RNA-seq (top); all individuals were distinct from those profiled with scRNA-seq. Flow cytometry analysis confirmed enrichment of classical monocytes (CD14+CD16-, bottom). (B) PCA of four patient and four control bulk transcriptomes from classical monocytes. (C) Significantly enriched gene sets between patient and control cohorts by GSEA. Dots are sized to denote significance (adjusted p-values); x-axis indicates NES. (D) Differentially expressed genes between patient and control. Dots are sized to denote significance (p-values).

### Monocyte dysregulation is heterogenous within and between ME/CFS patients

Our data identifies dysregulation of classical monocytes as a prominent feature of the immune system in ME/CFS. However, it is unknown whether this feature derives from consistent alterations across classical monocytes, or alternatively, is restricted to a subset of cells per individual. Similarly, bulk transcriptome profiling of classical monocytes from patients suggests more extensive variation between patients than between controls (Figure 3B), consistent with the possibility that patients possess a varied and heterogenous population of these cells, in contrast to a more consistent and homogenous population in controls. To explore these possibilities using the single-cell data for classical monocytes, we utilized a machine learning approach, positive unlabeled learning (Elkan & Noto, 2008), which accommodates mixed populations (Figure 4A). Starting with baseline case and control female samples, we optimized a classifier that labels each cell as diseased or normal (Figures 4B and S4A). The results indicated that ME/CFS patients possess a heterogenous population of classical monocytes, only some of which are diseased, with the remainder comparable to those in control individuals (Figures 4A and S4A). The percentage of cells classified as diseased within patients was variable (Figure 4C), but consistent for each individual when compared between the baseline and post-CPET samples (Figure 4D). We calculated the CH (Calinski-Harabasz) index applied within PCA space, as a metric of performance for our classifier in comparison with other sample metadata. The CH index analysis demonstrated that cells predicted as diseased are more highly related to one another than those partitioned by disease status, sex or the identity of an individual (Figure 4E), suggesting that the classifier is recovering latent signal within the data.

**Figure 4.**
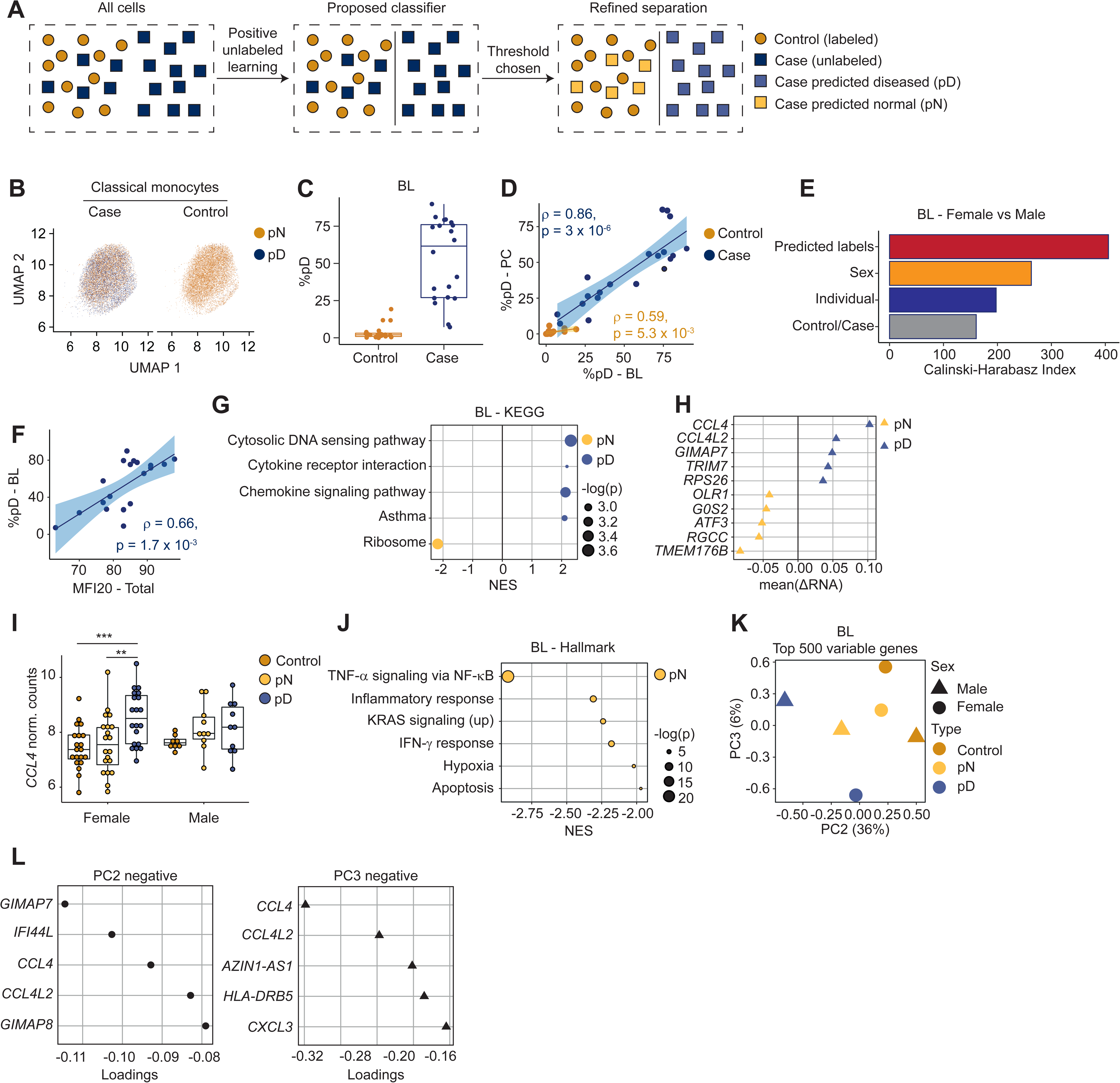
Heterogeneity in classical monocyte cells from ME/CFS patients. Data represent baseline female samples, unless described otherwise. (A) Schema describing positive unlabeled learning strategy to stratify single-cell transcriptomes from patients. (B) UMAP for classical monocyte cells, tiled and colored by predicted disease state. (C) Percentage of cells predicted as diseased (pD) per individual. (D) Correlation (Spearman) between percentage of pD cells per individual, compared between baseline and post-CPET. (E) Calinski-Harabasz index comparing performance of clustering across different stratifications of the single-cell dataset (y-axis). (F) Correlation (Spearman) between MFI-20 score and percentage of pD cells in cases. (G) Gene sets most differentially enriched between pD and predicted-normal (pN) cells from cases. Dots are color-coded to indicate enrichment in pD (blue) or pN (yellow) cells; sizes indicate corrected *P* values. (H) Genes (y-axis) most differentially expressed (x-axis) between pD and pN cells in an intra-sample paired analysis. (I) Expression of *CCL4* (y-axis) from individual samples, aggregating expression over pD and pN cells from cases and over all cells from controls (x-axis and color-coded). (J) Top 6 gene sets that are differentially enriched between pD and pN cells, based on GSEA of differential expression of genes between each subset of cells paired by sample. (K) Pseudobulk PCA from aggregated cells from control samples (yellow), pN cells from case samples (green), and pD cells from case samples (blue), partitioned also by sex. (L) Genes contributing to negative values for principal components 2 (top) and 3 (bottom) in panel K.

We examined correlations between the proportion of aberrant monocytes per individual and parameters that reflect patient health. Notably, we observed a significant correlation between the fraction of classical monocytes identified as diseased and female patients’ MFI-20 score (Smets et al., 1995; Multidimensional Fatigue Inventory-20) (Figure 4F), a comprehensive evaluation of fatigue. Similar correlations were also observed with other symptom metrics for females, including general health, SF-36 physical component scores (Nacul et al., 2011) and PEM severity (Figures S4D-F). We observed similar trends for the classifier trained with male samples (Figures S4B, C, and G-H). These observations indicate a relationship between symptoms ME/CFS patients experience and the fraction of diseased monocytes in circulation.

To examine the dysregulation of predicted diseased (pD) cells, we performed differential expression between patient cells predicted as normal (pN) versus diseased and used GSEA to examine patterns of differential expression. This analysis revealed clear differences between the gene expression programs (Figure 4G). Strikingly, the transcriptome profiles from pD cells compared to pN cells from patients exhibited changes in expression of pathways involving cytosolic DNA sensing, cytokine-cytokine receptor interaction, and chemokine signaling, which were similarly observed when comparing profiles of all patient cells to healthy controls in females (Figure S4I). In addition, ribosomal gene sets were also downregulated in pD cells in both females and males (Figures 4G and S4J), consistent with observations we made across multiple cell types (Figure 2C).

To minimize the impact of inter-individual variation in gene expression unrelated to disease state, and therefore more confidently identify genes dysregulated due to ME/CFS, we performed a paired analysis by individual of origin. Accordingly, we calculated the pseudobulk gene expression changes between pD and pN monocytes from the same ME/CFS individual, and then averaged these expression ratios across individuals. In this analysis, *CCL4* (C-C motif Chemokine ligands 4) exhibited the strongest change, with elevated expression in pD cells compared to pN cells within the same individuals (Fig. 4H). We also observed differences in *CCL4* expression levels when comparing profiles of pD and pN from patients to controls (Figure 4I). Elevated expression of *CCL4* in predicted-diseased cells was more prominent in females than in male patients (Figure 4I). Furthermore, *TMEM176B* (Transmembrane Protein 176B) was downregulated in pD cells compared to pN cells in ME/CFS individuals as well as control cells (Figures 4H and S4K). *TMEM176B* is involved in maintaining the immature state of dendritic cells (together with *TMEM176A*), which is anti-inflammatory (Condamine et al., 2010). Other genes upregulated in predicted diseased cells included *GIMAP7* and *TRIM7*, which may contribute to cell survival by suppressing apoptosis (Limoges et al., 2021) and promoting inflammation (Lu et al., 2019). The identities of genes downregulated in pD cells, such as *OLR1*, *G0S2*, *ATF3*, *RGCC,* and *TMEM176B* contain both pro-inflammatory and anti-inflammatory factors (Arslan et al., 2017; Kwok et al., 2020; Labzin et al., 2015; Okabe et al., 2022; Picotto et al., 2020). This observation may be attributed to the fact that the environment eliciting inflammatory responses is complex, with multiple ME/CFS specific changes in signaling occurring. For example, *ATF3*, upregulated in pD cells, is a regulator of interferon responses and able to suppress *CCL4* in animal models (Khuu et al., 2007; Labzin et al., 2015). We also performed GSEA using the paired analysis data (Figure 4J). The IFN-γ pathway was upregulated in pN cells, suggesting reduced inflammatory responses of pD cells through this pathway (as in Figure 3C).

Finally, we used PCA to visualize sex-specific pseudobulk transcriptomes generated from all control monocytes and from all patient cells partitioned by the classifier into predicted diseased and predicted normal. Transcriptomes derived from pD cells clustered away from control cells in PCA space and also from pN cells from the same patients (Figure 4K). In particular, PC1 reflected the sex of the individuals (Figure S4L), with PC2 and PC3 coincident with predicted disease state for males and females, respectively (Figure 4K). These observations suggest that the classifier separated the case cells into two groups: pD cells distinct from control cells, and pN cells that are less different than, but still show some deviation from, control cells. Analysis of loadings of the principal components (Figure 4L) show that *CCL4* (C-C motif Chemokine ligands 4) has a strong negative contribution to both PC2 and 3, indicating that dysregulation of this gene contributes strongly to discrimination between pD and pN cells in both males and females. Thus, within patients, a subset of classical monocytes exhibited high expression of cytokine receptor and chemokine signaling genes, in particular *CCL4*. Overall, these analyses demonstrated that classical monocytes populations are heterogenous within and variable across ME/CFS patients. Importantly, the classifier enabled us to identify predicted-diseased cells in ME/CFS patients and establish that these cells upregulate expression of specific cytokine receptor and chemokine signaling genes indicative of aberrant monocyte recruitment to tissue. Our observation that the percentage of diseased cells per individual correlate with metrics of disease severity is notable (Figures 4F and S4F).

### Signaling pathways impacting monocytes in ME/CFS

As monocytes in ME/CFS patients show significant dysregulation in pro-inflammatory chemokine and cytokine signaling pathways, they may be intrinsically biased towards this pre-migratory transcriptome, or alternatively, they may be responding differentially to changes in intercellular signaling in ME/CFS individuals. To assess whether intercellular signaling is altered in ME/CFS, we used CellChat (Vu et al., 2022) to analyze scRNA-seq data to model communication between cell types as a function of ligand and receptor expression levels corresponding to established signaling pathways (Jin et al., 2021) (Figure 5A). To remove potentially confounding factors such as sex, exercise and cluster size, we focused our analysis on the larger female-only cohort at baseline and down-sampled large clusters to 500 cells before calculating communication probabilities, an approach recommended for similar analyses (Andrijevic et al., 2022). Comparing the interactomes of patient and control cells, we found widespread evidence suggesting alterations in inferred communication to and from several cell types. In particular, monocyte signaling to certain T and NK cells is predicted to be elevated in ME/CFS patients, while also increasing signaling interactions among themselves (Figures 5B and S5A). These results suggest that monocyte dysregulation in ME/CFS, in part, may derive from alterations to established intercellular communication pathways.

**Figure 5.**
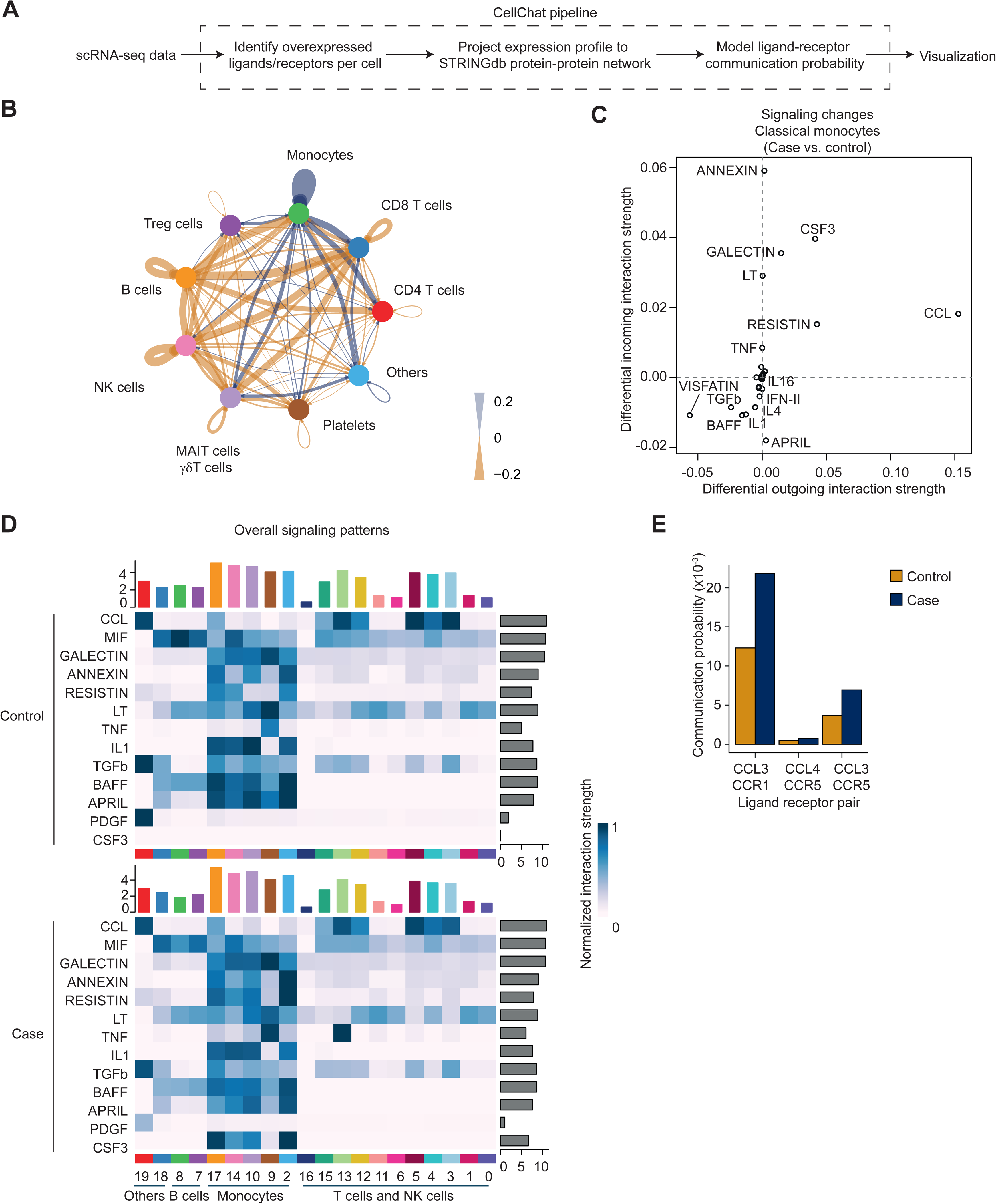
Intercellular signaling in the circulating ME/CFS immune system. (A) Schema depicting the CellChat strategy to detect cell-cell communication in scRNA-seq datasets. (B) Circle plot showing the differential number of interactions (case minus control), aggregating clusters of similar cell types. Blue indicates case cells exhibit more interactions than control cells; orange indicates control cells exhibit more interactions. (C) Scatter plot of differential incoming versus outgoing interaction strength in classical monocyte cells (cluster 2). Positive values indicate increased signaling strength in patients, and *vice versa*. (D) Heatmap of overall signaling for pathways dysregulated (y-axis) for classical monocytes receiving signaling from different cells (y-axis; cluster identifiers from Figure 1). Top bar plot indicates aggregate interaction strength of incoming signals; right bar plot indicates aggregate interaction strength of outgoing signals. (E) Bar plot of communication probabilities between specific ligand-receptor pairs in the CCL pathway, shown separately for case (blue) and control (orange) cells.

We next investigated the identities of the pathways that may contribute to alterations in the interactome of classical monocytes in ME/CFS. Among patients, we found evidence of increased signaling in pathways that regulate monocyte survival and localization to inflamed tissue (Galectin, Resistin, CSF3, CCL) (Hollmén et al., 2016; Hornig et al., 2015; Yıldırım et al., 2015), with the strongest dysregulation observed in genes associated with the C-C motif chemokine (CCL) pathway (Figures 5C-D and S5B), observations consistent with our earlier findings. To identify the specific ligand-receptor pairs contributing to overall changes in signaling in ME/CFS and with cell-type resolution, we compared the communication probabilities between the patient and healthy cohort, finding 782 upregulated and 1046 downregulated ligand-receptor pairs in patients, aggregated across all pairwise combinations of cell types (Supplementary File S7). In particular, for ligand-receptor pairs associated with the CCL pathway, we identified increased signaling from classical monocytes to platelets in ME/CFS patients, and this signal derived from elevated *CCL3*/*CCL5* expression in monocytes (Figure 5E). Monocyte-platelet interactions, especially in the form of monocyte-platelet aggregates, have previously been demonstrated to be a potent marker of inflammation (Passacquale et al., 2011; Stephen et al., 2013). Thus, classical monocytes in ME/CFS patients may exist in an inflammatory state, in part, due to altered cross-talk between platelets and monocytes.

### An abnormal platelet state is coincident with PEM in ME/CFS

Post-exertional malaise is a defining symptom of ME/CFS. To investigate whether PEM is associated with any changes in immune cells, we compared expression changes across different immune cells between baseline and 24 hours after CPET. Conventional analysis of differential gene expression comparing the average post-CPET expression to average baseline did not identify significant genes within cases or controls. Because our study design includes samples collected before and after exercise from the same individual, we could leverage a paired analysis to reduce inter-individual variation, which can compromise signal detection. For each cell-type (cluster), we calculated the expression ratio for each gene per individual in response to exercise, retaining only genes that met detection criteria. This approach generates a ΔRNA metric per individual, which we compared between the cohorts of patients and controls (Figure 6A). Tests for differential expression of individual genes again failed to reach significance. However, GSEA, which can detect more subtle but coordinated shifts in gene expression across a large number of genes, showed a marked number of enriched gene sets in platelets, with minimal signal or no signal in other cell types (Figure 6B).

**Figure 6.**
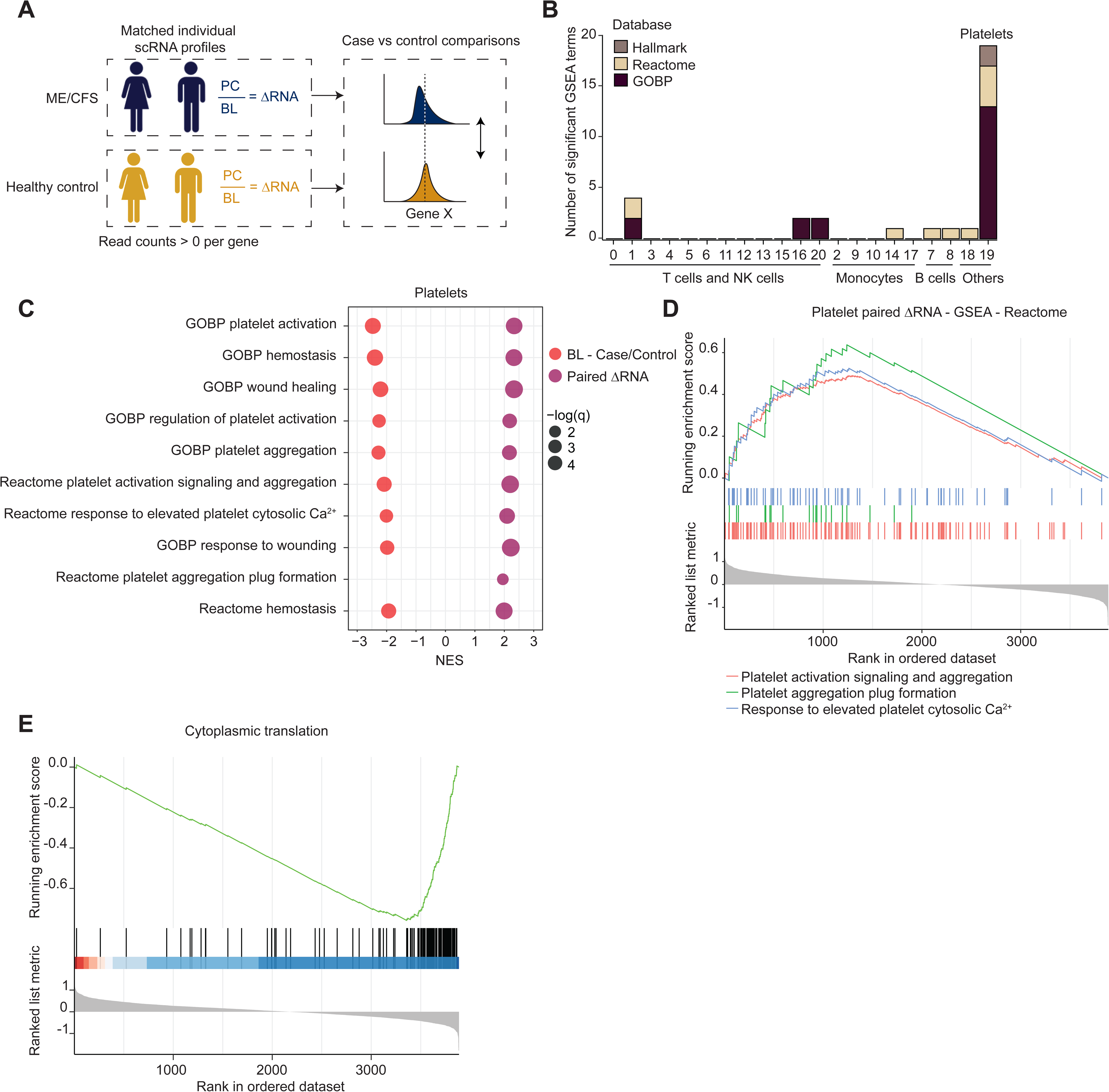
Aberrant platelet transcriptomes coincident with PEM in ME/CFS. (A) Schema depicting paired analysis (intra-individual expression) of gene expression altered by strenuous exercise in ME/CFS patients compared to controls. (B) Total number of significantly enriched gene sets across clusters (x-axis) in a paired analysis comparing case and control ΔRNA measurements with GSEA. (C) GSEA results for significantly enriched gene sets related to platelet function. GSEA analyses included the comparison of paired ΔRNA measurements between cases and controls (dark purple) as well as comparing group-averages at baseline (red) and post-CPET (none detected). (D) Enrichment plot depicting representative gene sets related to platelet function using the intra-individual paired approach. (E) Enrichment plot depicting a representative gene set related to translation using the intra-individual paired approach.

Gene sets related to platelet function were significantly enriched in the paired analysis of platelets, with positive enrichment in case ratios (post-CPET/baseline) compared to controls (Figures 6C and 6D). To examine the behavior of platelets at each timepoint, we repeated the GSEA analysis directly comparing ME/CFS patients and control cohorts at baseline and post-CPET. This analysis revealed reduced expression of platelet-function gene sets at baseline in the ME/CFS cohort compared to controls (Figure 6C). Interestingly, these enrichments were not evident in the post-CPET samples. These analyses revealed that the platelet cluster manifests a strong signature of dysregulation only prior to exercise in ME/CFS patients, an observation which suggests that strenuous exercise alters platelets in ME/CFS individuals.

Other notable gene sets in the paired GSEA analysis for platelets shared a high number of ribosomal protein genes (Figure 6E, Supplemental File S8). Again, this enrichment derived from differences in platelets at baseline in ME/CFS cases compared to controls, with no significant enrichment in the post-CPET cohorts. Therefore, the negative enrichment scores for the paired analysis reflects an increase in the detection of ribosomal proteins and translation machinery at baseline in ME/CFS cases. Notably, no other cell types exhibited a significant change due to exercise in the paired GSEA analysis (Figure 6B).

We note that aberrant platelet activation and fibrin amyloid microclots have recently been reported in patients with Long COVID (Pretorius et al., 2022) as well as in ME/CFS patients, though the microclot load was found to be less in the ME/CFS cohort (Nunes et al., 2022). These studies correspond most closely to the baseline state assessed here, and our results also suggest that dysregulation, as judged by transcriptome analysis, occurs at baseline. Nevertheless, it is clear that the CPET induces a marked change in the transcriptomes of platelets in ME/CFS individuals.

## DISCUSSION

This study provides a new and important resource to investigate immune dysregulation in ME/CFS. Here, because classical monocytes manifested the strongest signal of dysregulation in ME/CFS, we focused on exploring changes in their gene expression program, as a novel and potentially important aspect of the disease. However, alterations to additional immune cells are clearly also an important aspect of ME/CFS, with the strongest signal observed in CD4+ T cell subsets within our data, including in antigen-inexperienced naïve CD4+ T cells. Patterns of dysregulation limited to a very small subset of cells, such as in adaptive immune cells specific to a particular antigen (or antigens), are difficult to detect in scRNA-seq data. Nevertheless, it is worth noting that we observe changes in gene expression programs in clonally diverse antigen-experienced CD4+ T cells and in other such adaptive cell populations, an observation consistent with dysregulation of adaptive immune cells occurring via cytokine-mediated bystander regulation (Shim et al., 2022) rather than via T-cell receptor specific interactions. Nevertheless, it is clear that the largest alterations in ME/CFS PBMCs detectable by scRNA-seq are found in monocytes.

Multiple lines of evidence implicate dysregulation of classical monocytes in ME/CFS. In particular, our analysis discovered upregulation of chemokine/cytokine pathway genes in patient-derived monocytes as well as a clear correlation between the proportion of predicted-diseased monocytes and multiple metrics of disease severity. Future studies investigating macrophages from tissue biopsies isolated from ME/CFS patients will be important in this regard. CellChat analyses also identified striking alterations in intercellular communication within the immune system of ME/CFS. In contrast to healthy controls, where a diverse network of intercellular interactions is predicted, monocytes are predicted to contribute additional overall information exchange in ME/CFS. This observation may be a consequence of a prolonged exposure to an inflamed environment in ME/CFS patients, which can alter cellular metabolism and functions (Lacourt et al., 2018). We also detected conflicting pathways related to immune response as upregulated in classical monocytes, specifically, both pro-inflammatory and anti-inflammatory responses. For example, although we detected an upregulation of the pro-inflammatory IL-1β pathway, the anti-inflammatory IL-10 pathway was also elevated within ME/CFS cases. These observations call for more extensive characterization of classical monocytes in ME/CFS to identify the overall response of such cells in an abnormal environment. Future studies that integrate plasma cytokine analysis with monocyte state in a common cohort of patients (and controls) will be particularly valuable.

How might aberrant monocyte activation contribute to symptoms experienced by ME/CFS patients? Monocytes express multiple types of chemokine receptors, which in response to different chemokines, direct monocytes to a variety of tissues. Here, we observed increased expression of *CCL3* and *CCL4*, which have been shown to direct monocyte homing to joints and adipose tissues in osteoarthritis and adipose tissue in obesity (Sindhu et al., 2019; Zhao et al., 2020). Thus, our observations suggest that ME/CFS patients experience continual improper recruitment of monocytes to one or more tissues. If this hypothesis is correct, it will be important to examine the balance of pro- and anti-inflammatory macrophages in tissues from patients, and whether the macrophages themselves are altered.

A previous study identified aberrant platelet activity in ME/CFS (Nunes et al., 2022), although another study, using different methods, did not observe alterations in platelets (Kennedy et al., 2004). There is clear evidence of aberrant platelet activation in Long COVID (Pretorius et al., 2021). Our analysis suggests an important refinement to the hypothesis linking platelet activation to ME/CFS (and perhaps Long COVID): although aberrant activation may be present at baseline, it appears that the platelet population in circulation undergoes a substantial change in response to strenuous exercise (CPET, within 24 hours), and therefore a clear association with PEM. Our data suggests a model in which the population of circulating platelets in patients shifts within 24 hours following strenuous exercise. At baseline, the patient platelet transcriptome is biased towards a lower expression program of genes important in platelet activation. Following exercise, the platelet transcriptome in patients looks relatively normal, indicating either a loss of platelets harboring defective transcriptomes or a large infusion of new platelets. We envision two models to explain our observations relating to platelet dysregulation in ME/CFS and the alteration in their gene expression post-CPET. First, perhaps a subset of platelets in patients exist in a state that renders them susceptible to activation, and strenuous exercise induces microclot formation, removing them from circulation; thus, post-CPET, only normal platelets remain. The second model envisions that exercise induces an influx of normal platelets, perhaps in combination with clearance (or clot formation) of older platelets.

Platelets, lacking a nucleus, have unconventional transcriptomes. Upon their release into circulation, platelets inherit the transcriptome and proteomes of their megakaryocytes of origin without introduction of any new transcripts. Therefore, the platelet transcriptome degrades without replenishment as platelets circulate in the body. The rate of RNA degradation *in vitro* shows a fast degradation with half of the RNA lost after six hours and almost all (98%) at 24 hours (Angénieux et al., 2016). Selective degradation of the platelet transcriptome has also been reported, where transcripts encoding the translational machinery degrade slower than others (Mills et al., 2017). Our analysis shows that platelets at baseline in ME/CFS subjects possess a transcriptome indicative of older platelets, with a pronounced reduction in transcript levels for genes essential to platelet activation and function and an increase in ribosomal protein genes and other genes relevant to translation. However, the exercise challenge erases such signals, which is a striking phenomenon. In healthy individuals, acute exercise has been reported to activate platelets (Heber & Volf, 2015) and upregulate both pro-inflammatory and anti-inflammatory cytokines (Docherty et al., 2022), but prior studies have not explored the effect of exercise on platelet function in ME/CFS.

Distinct from monocyte dysregulation, one of the most prominent features of transcriptome dysregulation in ME/CFS is repression of translational machinery and ribosomal protein genes, which we observed across multiple cell types. Regulation of such genes is complex, involving multiple pathways including p53 and TOR (Kang et al., 2021), although in general, the repression we observe suggests that multiple immune cells exist in a more quiescent or less proliferative state than normal. For example, normal activation of CD8+ T cells requires increased translation (Araki et al., 2017). It is worth noting that both CD4+ and CD8+ T cells in ME/CFS exhibit reduced glycolysis (Mandarano et al., 2020) and NK cells in patients are also known to have impaired cytotoxic activity (Eaton-Fitch et al., 2019). Future studies could be designed to systematically examine these questions, first testing for correlations between these molecular changes, and if such correlations exist, seeking a mechanistic understanding of them.

Importantly, ME/CFS and Long COVID, together with other post-viral diseases, have been suggested to share common molecular alterations (Tate et al., 2022). Nevertheless, at present this suggestion is, largely, based on an overlapping (although not identical) set of symptoms, rather than molecular or cellular data. Our high-resolution data, describing the circulating immune system in ME/CFS, will be an ideal comparison set for future studies of Long COVID, with the potential to identify both congruent and divergent aspects of immune function in ME/CFS and Long COVID. In this regard, future studies will determine whether classical monocyte dysregulation is also the most prominent signal in the circulating immune system of Long COVID patients, and more importantly, whether the genes and gene sets impacted in ME/CFS are also observed in monocytes from Long COVID patients.

## Acknowledgements

This study was supported by NIH grant U54NS105541 to M.R.H., A.G. and D.C.S., and by UL1 TR 002384 from the National Center for Advancing Translational Sciences (NCATS) of the National Institutes of Health (NIH). We thank the Cornell Biotechnology Resource Center (BRC) Genomics Facility, Flow Cytometry Facility, and Transcriptional Regulation and Gene Expression Facility (TREx) for providing excellent services. Thanks to Dr. John Chia for invaluable support in recruiting and screening ME/CFS individuals and controls. We would like to send our heartfelt thanks to the ME/CFS subjects who participated in this study.

## Author Contributions

Conceptualization A.G.; Methodology and Software F.A., H.Z., J.K.G; Validation L.T.V. and E.A.F.; Formal Analysis F.A., H.Z., D.S.H.I., Y.K., W.C., P.R.M. and J.K.G.; Investigation F.A., L.T.V., H.Z., E.A.F., D.S.H.I., Y.K., W.C., P.R.M. and J.K.G.; Resources C.J.F., G.E.M., S.M.L., J.K.C, B.A.K., J.S., M.R.H., X.M. and D.C.S.; Data Curation A.F., J.K.G.; Writing – Original Draft A.G.; Review and Editing F.A., L.T.V., H.Z., D.S.H.I., E.A.F., M.R.H., J.K.G. and A.G.; Visualization F.A., L.T.V., H.Z., D.S.H.I. and J.K.G; Supervision and Project Administration A.G. and J.K.G.; Funding Acquisition A.G., M.R.H. and D.C.S.

## METHODS

### RESOURCE AVAILABILITY

#### Lead contact

- Further information and requests for resources and reagents should be directed to and will be fulfilled by the lead contact, Andrew Grimson.

#### Materials availability

- This study did not generate new unique reagents.

#### Data and code availability

- De-identified human single-cell and bulk RNA-seq data have been deposited at GEO.
- Any additional information required to reanalyze the data reported in this paper is available from the lead contact upon request.

### EXPERIMENTAL MODEL AND SUBJECT DETAILS

#### Human Subjects

The human subjects research described in this publication was approved by the Weill Cornell Medical College and Ithaca College Institutional Review Boards and participants provided written informed consent prior to participation as previously described (Germain et al., 2022). Participants at 3 sites donated PBMCs and demographic information as part of the Cornell ME/CFS Collaborative Research Center. The specific cohort in this study was selected using age, BMI, sex, and peak VO_2_ from cardiopulmonary exercise tests (CPETs). Age and BMI ranges were matched to minimize confounding factors between cases and controls (Figure 1B). Sex was considered to ensure that the cohort matched disease prevalence where considerably more females than males report having ME/CFS (Bakken et al., 2014; Jason et al., 1999). Cases were selected that demonstrated a considerable decrease in peak VO_2_ over the course of 2 CPETs conducted as part of the larger study (Germain et al., 2022) (Figure 1B), as an objective basis for physiological dysfunction within the ME/CFS participants (Keller et al., 2014).

The Multidimensional Fatigue Inventory (MFI-20) (Smets et al., 1995) was used to assess the level of fatigue in cases. The SF-36v2® Health Survey (John E. Ware, 2007) was used to compare general health and quality of life between cases and controls. A modified version of the Chronic Fatigue Syndrome severity score (Baraniuk et al., 2013) was used to measure post-exertional malaise (PEM). The severity score measured PEM on a 0-10 scale over the past month. The MFI-20, SF-36v2, and PEM severity score are all self-reported. The MFI-20, SF-36v2, and past month-PEM severity were completed before visiting the test site to perform the CPETs. Current PEM severity was then serially measured at the test site prior to CPET and then every two days post CPET for at least ten days to measure recovery.

In order to preserve anonymity in the public data repository, the subjects’ ages at the time of sample collection are coded by age bin (1: >= 18 to <= 35 years; 2: > 35 to <= 45 years; 3: > 45 to <= 55 years; 4: > 55 to <= 70 years).

## METHOD DETAILS

### PBMC isolation

ME/CFS cases and healthy sedentary controls participated in a larger study conducted by the Cornell ME/CFS Collaborative Research Center as previously described (Germain et al., 2022). This study utilized PBMCs processed from whole blood collected from each participant over the course of two days, separated by a cardiopulmonary exercise test (CPET, Figure 1A).

Whole blood was collected in EDTA tubes and centrifuged on the day of collection for 5 minutes at 500 rcf prior to removing the plasma fraction. The remaining sample was diluted with equal parts PBS and transferred to a SeptMate^™^ Tube (Stemcell Technologies) containing Histopaque^®^-1077 (Sigma-Aldrich). SeptMate^™^ Tubes were then centrifuged for 10 minutes at 1,200 rcf and the buffy coat layer was transferred to a new tube. After two washes with PBS (first wash was centrifuged for 10 minutes at 120 rcf for without brake; the second was centrifuged for 8 minutes at 300 rcf), the resulting pellet was resuspended in PBMC storage media (60% RPMI 1640, 30% heat inactivated FBS, and 10% DMSO), counted, and divided into aliquots of ~1 - 10 million cells per mL. PBMC aliquots were promptly transferred to −80 °C for slow freeze down in a Mr. Frosty™ Freezing Container (Thermo Scientific). After 24 hours, the PBMCs were transferred to liquid nitrogen for long-term storage.

### Single-cell gene expression profiling

Samples were co-processed for single-cell RNAseq (scRNAseq) in batches of 4-8 PBMC aliquots, such that each batch contained paired baseline and post-CPET samples from the same individual and a mix of cases and controls. PBMCs were prepared for scRNAseq using the 10x Genomics Demonstrated Protocol: Fresh Frozen Human Peripheral Blood Mononuclear Cells for Single Cell RNA Sequencing (CG00039) as a guide. Briefly, vials were rapidly thawed in a 37°C water bath and 500µL was transferred to a 50ml centrifuge tube using a wide-bore tip. Cells were serially diluted 1:1 with RPMI 1640 (Gibco # 11875093) plus 10% heat inactivated FBS (Gibco #10438026) in one-minute increments until the total volume reached 32 ml. Cells were centrifuged in a swinging bucket rotor at 300 rcf for 5 minutes at room temperature. After discarding the majority of the supernatant, cells were gently resuspended in the residual volume with a regular-bore tip to achieve single cell suspension, transferred to a 15ml centrifuge tube and the volume brought up to 10ml with RPMI 160 plus 10% heat inactivated FBS. Cells were centrifuged again at 300 rcf for 5 minutes at room temperature, and the pellet was gently resuspended in 1 ml of 1x PBS (BioWhittaker #17-516F) plus 0.04% BSA (Invitrogen #AM2616) with a wide-bore tip. The cells were transferred to a 2 ml microfuge tube, centrifuged at 300 rcf for 5 minutes in a swinging bucket centrifuge at room temperature, and resuspended in 1x PBS plus 0.04% BSA to a specified cell concentration to load ~8,000 cells onto the 10x Chromium chip for a target capture of 5,000 cells. Cells were counted on a TC-20 cell counter (Bio-Rad) multiple times to track total and live cell counts. In total, 120 PBMC samples were processed in 18 batches and only two paired samples from a control individual failed the minimum viability tests and were excluded from the study.

Viable cells were submitted to the Cornell BRC Genomics Facility for processing with the Chromium Single Cell 3L v3 kit (CG000183 Rev A and Rev B, 10x Genomics). The Facility prepared a total of 120 single-cell RNAseq libraries; all underwent quality checks for size distribution on a Fragment Analyzer 5200(Agilent) and molarity on a QX100 Digital Droplet PCR Machine (Bio-Rad). Libraries were sequenced in an initial run to generate preliminary data quality metrics, and additional sequencing depth was generated as required to meet the target coverage for each sample. The libraries generated in batches 1-13 were primarily sequenced on a NextSeq500 (R1:28bp, R2:55bp) and the libraries from batches 14-18 were sequenced on a HiSeq2000 followed by a NovaSeq6000 (both PE 2×150bp). A final sequencing run on a NextSeq2000 brought all samples to the minimum target depth (see below).

### Monocyte isolation and profiling

#### Magnetic enrichment

Samples from individuals in the larger Cornell Center study that were not included in the scRNA-seq cohort were selected for monocyte profiling. Specifically, we used PBMCs from eight post-CPET females (4 cases with low SF36v2 PCS scores and 4 controls, matched by age and BMI). Vials containing cryopreserved PBMCs were rapidly thawed in a 37°C water bath and serially diluted 1:1 with RPMI 1640 plus 10% heat inactivated FBS in one-minute increments until the total volume reached 32 mL. Cells were centrifuged in a swinging bucket rotor at 300 rcf for 5 minutes at room temperature and the pellet was gently resuspended in 10ml MACS buffer (Miltenyi Biotech Cat# 130-091-376). After two washes in MACS buffer, centrifuging at 300 rcf for 5 minutes at room temperature, the pellet was resuspended in 1 ml of MACS buffer and cells were counted with a TC-20 cell counter (Bio-Rad). Monocytes were initially purified with the Classical Monocyte Isolation kit, human (Miltenyi Biotech Cat#130-117-337), then additionally enriched for CD14+ cells using the CD14 MicroBeads (Miltenyi Biotec Cat#130-050-201). After counting, 20,000 – 140,000 cells were removed to 250µl of MACs buffer and 750µl Trizol LS (ThermoFisher) added to lyse cells for bulk RNAseq profiling. Trizol lysates were frozen at −80 °C prior to RNA extraction and submitted to the Cornell Transcriptional Regulation and Expression (TREx) Facility for RNA extraction and RNAseq. Remaining cells were immediately analyzed with flow cytometry.

#### Flow cytometry

Human TruStain FcX™ Fc Receptor Blocking Solution (BioLegend Cat#422302) was added according to the number of cells, mixed well and incubated for 10 minutes at room temperature. Cells were divided into separate microfuge tubes for staining with fluorescently-labeled antibodies (CD3-FITC, CD45-PerCP, CD16-PE, and CD14-APC-CyC7 from BioLegend) for flow cytometry and incubated on ice for 30 minutes. 1 ml of FACS buffer (heat inactivated FBS, 0.5M EDTA pH8.0 1x PBS) was added to the cells, mixed well and centrifuged at 300 rcf for 5 minutes at 4°C. The supernatant was removed and the cells were washed once more. The final cell pellet was resuspended in 100µl of FACS buffer and kept on ice in the dark. Sytox blue (0.2µl/ 100µl or less) was added directly before flow cytometry analysis and analyzed on a Thermo Fisher Attune NxT at the Cornell BRC Flow Cytometry Facility.

#### Bulk RNAseq

At the TREx Facility, RNA was extracted from Trizol following the manufacturer’s protocol with the following exceptions: the aqueous fraction was re-extracted with an equal volume of chloroform in Phase Lock Gel Heavy tubes (QuantaBio) and 2 µl GlycoBlue (Thermo) was added prior to precipitation to improve RNA recovery. Total RNA samples were quantified with the Qubit HS RNA assay (Thermo) and integrity assessed on a Fragment Analyzer (Agilent) to confirm RQN values ≥ 7. Bulk polyA+ RNAseq libraries were generated from 25ng total RNA with the NEBNext Ultra II Directional RNA kit (New England Biolabs). Libraries were quantified with a Qubit HS DNA assay (Thermo) and sequenced on a NovaSeq6000 (Illumina) at Novogene to generate a minimum of 20M PE 2×150bp reads per sample.

## QUANTIFICATION AND STATISTICAL ANALYSIS

### Single-cell RNAseq

#### Data processing

Fastq files were generated with cellranger mkfastq (10x Genomics) by the sequencing facility (BRC Genomics Facility or Novogene). Raw count tables were generated with cellranger count v6 (10x Genomics) [cellranger count --id=ID --transcriptome=/path/to/refdata-gex-GRCh38-2020-A/ --fastqs=/path/to/directories --sample=list --expect-cells=5000 --r1-length=28 --r2-length=55 –nosecondary]. Because sequencing read lengths from different instruments varied, and this was observed to contribute to bias in the count tables, reads were trimmed to match the minimum length across the dataset (R1 = 28nt, R2 = 55nt).

#### Single-cell integration and clustering

Count tables from all samples were imported into R to analyze with Seurat (v4.1.0) (Hao et al., 2021). Initial filtering removed cells that did not meet minimum quality criteria (nFeature_RNA > 500 & nFeature_RNA < 5000 & percent.mt < 30 & log10GenesPerUMI > 0.80). A pair of samples from the same control individual were discarded due to an excess of counts for mitochondrial genes indicating a sample quality problem, leaving a total of 116 samples in the final dataset. Normalization of UMI counts for each library was performed using the SCTransform function [SCTransform(sobj, method = “glmGamPoi”, vars.to.regress = “percent.mt”, return.only.var.genes = FALSE)]. Samples were integrated with the R script RunHarmony (Korsunsky et al., 2019) [wrapper for RunUMAP(sobj, reduction = “harmony”, dims = 1:50)). Clustering with Seurat [FindNeighbors(sobj, reduction = “harmony”, dims = 1:50) %>% FindClusters(resolution = 0.6] generated a total of 29 clusters. Cell doublets within each sample were determined with DoubletFinder (v2.0.3) (McGinnis et al., 2019) and removed from the dataset. The FindMarkers() function in Seurat was used to determine marker genes between clusters and genes that distinguish case and control cells within each cluster.

#### Pseudobulk analysis

Raw pseudobulk counts were extracted from the Seurat object as the sum of counts per sample, per cluster. Normalization of pseudobulk matrices with DEseq2 (v1.30.0) (Love et al., 2014) generated normalized counts for downstream analyses, and was rerun as the cell assignments or cohorts were altered for different analyses. DEseq2 was used to detect differentially expressed genes between groups, using ‘minReplicatesForReplace = Inf’ to reduce the contribution from spurious outliers.

The log_2_-fold change calculation from DEseq2 was used for gene set enrichment analysis (using R packages clusterProfiler (default parameters: v3.18.1) (Yu et al., 2012) or fgsea (Korotkevich et al., 2021) (minimal gene set size 5, maximum gene set size 2000, eps 0: v1.20.0) to run the GSEA (Mootha et al., 2003; Subramanian et al., 2005) algorithm, after filtering out genes with low coverage. For example, the GSEA analysis for clusters with more than 10,000 cells was filtered to retain only the top quartile of genes based on the median normalized counts across all samples. The Hallmark, C2:CP and C5 catalogs from the MSigDB database (Subramanian et al., 2005) were used for enrichment tests.

For the paired analysis controlling for the individual of origin, a ratio was calculated for each gene and each individual when normalized counts were available for both measurements and only for individuals with at least 4 cells contributing to pseudobulk counts per timepoint. Normalized counts were floored to 1 to reduce the contribution from poorly detected genes. GSEA analysis used the log_2_-fold change metric calculated from the geometric mean of case and control ratios per individual for genes with at least 6 ratios per group, and the Wilcoxon rank-sum test was used to compare groups.

Scores were calculated per sample (or per ratio) with the R package singscore (Foroutan et al., 2018), using a unique list of genes derived from the GSEA leading edge genes from gene sets related to chemokine/cytokine signaling (listed in Figure 2D), and the Wilcoxon rank-sum test was used to compare groups.

#### Cell demographics analysis

Cell counts per cluster per individual were normalized to the overall cell counts per individual library. We performed Wilcoxon rank-sum tests for the normalized cell counts in the healthy versus disease cohort at baseline and post-exercise respectively, as well as for normalized cell counts at baseline versus post-exercise separately for cases and controls.

#### Machine Learning classifiers

Positive unlabeled learning was performed based on publicly available methods and codes with the scRNA-seq dataset as input (Alon Agmon, 2020/2022; Elkan & Noto, 2008). Briefly, the algorithm took the normalized single cell gene expression matrix of cluster 2 (classical monocytes), using data from either females or males at baseline or post-CPET (data slot of the RNA assay in the Seurat object) as input. Cells from control individuals are labeled as positive, and cells from case individuals are unlabeled. Twenty percent of labeled cells (i.e., cells from control individuals) were held out, and the remaining 80% labeled cells together with all unlabeled cells (i.e., cells from case individuals) were used to train an XGB classifier (using Python package XGBoost (Chen & Guestrin, 2016), with default parameters) separating labeled and unlabeled cells. Next, the reserved labeled cells were projected onto the labeled/unlabeled classifier to estimate the probability of cells being labeled if they are positive (control-like). In the following step, all labeled and unlabeled cells were projected onto the labeled/unlabeled classifier. Finally, based on the theorem of conditional probability, the probability of unlabeled cells being positive can be estimated by the probability of the cells being labeled divided by the probability of cells being positive when labeled. Predicted probabilities were averaged across 24 iterations.

To select the probability threshold that determines if a cell is predicted as healthy or diseased, the Calinski-Harabasz Index of predicted diseased and predicted control cells was calculated based on the top 50 principal components of the single cell expression matrix across thresholds 0.1-0.9 (Figure S4A). The threshold with the highest Calinski-Harabasz Index was selected (0.4 for classical monocytes in females at baseline). Cells with probabilities higher than the threshold are considered as predicted control, and cells with probabilities lower than the threshold are considered as predicted diseased.

To correlate the classifier performance with other metrics, the percentage of cells predicted as diseased were calculated for each individual. Spearman correlation was calculated by R package ggpubr v0.4.0 between percentage of cells predicted diseased at baseline compared to post-CPET in cases and controls, or between percentage predicted-diseased cells and demographic metrics (MFI-20 total score, general health score, SF-36 physical component score, and PEM maximum change) across individuals. PEM maximum change was calculated as the largest delta for the PEM symptom severity scores (from baseline to each of the survey timepoints post-CPET).

To evaluate classifier performance, Calinski-Harabasz Indexes of cells partitioned by predictions and other sample metadata (sex, case/control, individual of origin) were calculated based on top 50 principal components of the partitioned single cell expression matrix in PCA space, generated with RunPCA() from Seurat.

Differential expression between cells predicted diseased and control was performed by FindMarkers function in Seurat. All genes are included. GSEA on log_2_-fold changes of genes between predicted diseased and predicted control groups of cells was performed using R package fgsea v1.20.0 (Korotkevich et al., 2021), with minimal gene set size = 5, maximum gene set size = 2000 and eps = 0. The Hallmark gene sets and KEGG subset from the C2 catalog in MSigDB database (Subramanian et al., 2005) were used for enrichment tests.

Paired analysis was performed by calculating the log_2_-fold changes between predicted-diseased and predicted-normal cells for each case individual. The ratios per genes were then averaged across case individuals and sorted based on differential expression levels. Pseudobulk gene expression was calculated by aggregating gene expression counts of cells partitioned by the predicted labels. Vst function in DESeq2 v1.34.0 (Love et al., 2014) was used to normalize the pseudobulk counts. PCA was performed on normalized pseudobulk counts using the R package prcomp and the loadings were then extracted from rotation of the PCA result.

#### Inference of cell-cell communication

CellChat (v1.1.3) (Vu et al., 2022) was used to infer cell-cell communication probabilities and identify signaling changes across healthy and diseased cell populations. We down-sampled each cluster to 500 cells from females, at baseline only, for a balanced comparison. We discarded clusters with fewer than 500 cells. Briefly, we first identified, for each cluster, differentially over-expressed ligands, receptors and cofactors in the human CellChatDB database (Jin et al., 2021), then their average expression values were used to calculate communication probabilities between all cell groups. Interaction strength along specific intercellular signaling pathways was calculated by summarizing the communication probabilities of associated ligand-receptor pairs across all clusters. Comparison of healthy and diseased signaling networks was performed as described previously (Vu et al., 2022), with statistical significance of changes in communication probabilities determined by Wilcoxon rank-sum test and using only cells collected from females at baseline.

### Flow cytometry analysis

Raw compensated flow cytometry files (.fcs) were analyzed using Flowjo (v10.8.1). Cells were first gated by size and granularity. Single cells were selected by looking at forward scatter area and height and side scatter area and height. Live cells were selected by gating on Sytox blue negative events. Bulk monocytes were selected as CD45+CD3- cells. Monocyte subsets were analyzed by investigating surface expression of CD14 and CD16. Classical monocytes (CD14+CD16-) were gated based on fluorescent-minus-one (FMO) controls.

### Bulk RNAseq

Fastq files were trimmed to remove 3′ low quality and adaptor sequences with TrimGalore (v0.6) (*Babraham Bioinformatics - Trim Galore!*, n.d.), a wrapper for cutadapt (Martin, 2011) and fastQC (*Babraham Bioinformatics - FastQC A Quality Control Tool for High Throughput Sequence Data*, n.d.), retaining reads ≥ 50bp. Trimmed reads were mapped to the reference genome (GRCh38 with Ensembl gene annotations) with STAR (v2.7) [--outSAMstrandField intronMotif, --outFilterIntronMotifs RemoveNoncanonical, --outSAMtype BAM SortedByCoordinate, --quantMode GeneCounts], which outputs a count table of reads per gene for each sample. DESeq2 (Love et al., 2014) was used to normalize raw counts, generate PCA (normalized by rlog) and MA plots, and analyze differential expression, using genes with more than 10 read counts in more than or equal to 3 libraries. The log_2_-fold change values from DEseq2 were used for GSEA analysis with R package fgsea v1.20.0 (Korotkevich et al., 2021) as described above, with minimal gene set size = 5, maximum gene set size = 2000, eps = 0 and gseaParam = 0. Hallmark, C2:KEGG and C2:Reactome gene sets from the MSigDB database (Subramanian et al., 2005) were used for enrichment tests.

## Supplemental figure legends

**Supplemental Figure 1. Single cell transcriptomics of the ME/CFS immune system**

Related to Figure 1

(A) Violin plot showing quality control metrics (nCount_RNA, nFeature_RNA, percent-MT, S.Score, G2M.Score) for each batch of samples (x-axis) processed on the 10x Genomics Chromium instrument. (B) Violin plot showing quality control metrics (nCount_RNA, nFeature_RNA, percent-MT, S.Score, G2M.Score) for each cluster (x-axis). (C) UMAP split by condition (Case-Baseline, Case-postCPET, Control-Baseline, Control-postCPET), showing representation of all clusters in each condition. (D) Relative cell numbers between cohorts across all clusters except clusters 4, 13, and 15 (see Figure 1F).

**Supplemental Figure 2. Dysregulation of immune cells in ME/CFS**

Related to Figure 2

(A) PCA plots for pseudobulk analysis of gene expression for cluster 2 (classical monocytes, top) and cluster 19 (platelets, bottom). Left panels are colored by sex, center panels are colored by condition (Case-Baseline, Case-postCPET, Control-Baseline, Control-postCPET), and right panels are colored by batch (Chromium processing). In most clusters and as shown for cluster 2, sex explains the first principal component of variation in the gene expression profiles. Cluster 19 is unique in not showing a strong sex bias. (B) GSEA results for comparisons of case versus control cohorts at baseline or post-CPET for clusters 9 (top) and 10 (bottom) for the same gene sets shown in Figure 2D, when the result is statistically significant (q-value < 0.05). Dots are sized to denote significance and colored to indicate the timepoint for the comparison of case versus control (Baseline or post-CPET); x-axis indicates normalized enrichment score (NES). (C) Single-sample scores for pseudobulk profiles for clusters 9 (top) and 10 (bottom) generated using the same list of genes as in Figure 2E (leading-edge genes from Figure 2D). * p-value < 0.05. (D) Correlation (Spearman) for single-sample scores for each individual, comparing baseline (x-axis) to post-CPET (y-axis).

**Supplemental Figure 3. Complete transcriptomics of classical monocytes**

Related to Figure 3

(A) MA plot showing average expression of genes (x-axis, DEseq2 baseMean) and the log_2_-fold change between case and control groups (y-axis). Dots are color-coded to indicate statistically significant differential expression and the group with higher relative expression (case, control; blue, yellow, respectively) at p-adjusted < 0.05. (B) Differential expression of genes associated with monocyte migration and differentiation for classical monocytes collected as in Figure 3A between case and control groups. Dots are sized to denote significance (p-values) and colored to reflect the group with higher relative expression (case, control; blue, yellow, respectively); x-axis indicates log_2_-fold change (case/control).

**Supplemental Figure 4. Heterogeneity in classical monocyte cells from ME/CFS patients**

Related to Figure 4

(A) Calinski-Harabasz Index comparing performance of classification for using different cutoffs for predictions, using female classical monocytes at baseline. The threshold with the maximum CH Index value for the classifier was selected (0.4). (B) Performance of a classifier for male cells: percentage of cells predicted as diseased (pD) per sample; males only, examined at baseline. (C) Correlation (Spearman) between percentage of pD cells per sample, comparing paired baseline and post-CPET values per individual; males only. (D) Correlation (Spearman) between general health score and percentage of pD cells per sample; females only, examined at baseline. (E) Correlation (Spearman) between SF-36 physical component score and percentage of pD cells per sample; females only, examined at baseline. (F) Correlation (Spearman) between maximum change in PEM symptom severity and percentage of pD cells per sample; females only, examined at baseline. (G) Correlation (Spearman) between MFI-20 total score and percentage of pD cells per sample; males only, examined at baseline. (H) Correlation (Spearman) between general health score and percentage of pD cells per sample; males only, examined at baseline. (I) Top 5 gene sets differentially enriched in GSEA comparing average log_2_-fold change (FindMarkers) in case and control cells from females at baseline. Dots are color-coded to indicate enrichment in case (blue) or control (yellow) cells and sized to indicate corrected P values. (J) Top 5 gene sets differentially enriched in GSEA comparing predicted normal (pN) and pD cells from males at baseline. Dots are color-coded to indicate enrichment in pN (yellow) cells and sized to indicate corrected P values. (K) Expression of *TMEM176B* (y-axis) across indicated groups (X-axis) per sample, aggregating expression of cells from controls (yellow), pN cells from cases (green), and pD cells from cases (blue), partitioned by sex, all at baseline. (L) PCA (principal components 1 and 2) of pseudobulk profiles from aggregated subsets of cells from controls (yellow), pN cells from cases (green), and pD cells from cases (blue), partitioned by sex, all at baseline.

**Supplemental Figure 5. Intercellular signaling in the circulating ME/CFS immune system**

Related to Figure 5

(A) Heatmap of differential interaction strengths between cell types at baseline, from female samples following down-sampling. Top bar plot indicates aggregate interaction strength of incoming signals to indicated clusters (X-axis); right bar plot indicates aggregate interaction strength of outgoing signals from indicated clusters (Y-axis). Positive values (blue) indicate increased signaling strength in cells from ME/CFS patients compared to controls; negative values (orange) indicate decreased signaling strength. (B) Violin plots of log-normalized expression levels for genes annotated under the CCL pathway in CellChatDB, per cluster, showing female cells at baseline in the control (orange) and ME/CFS (blue) cohorts.

## Supplemental Files

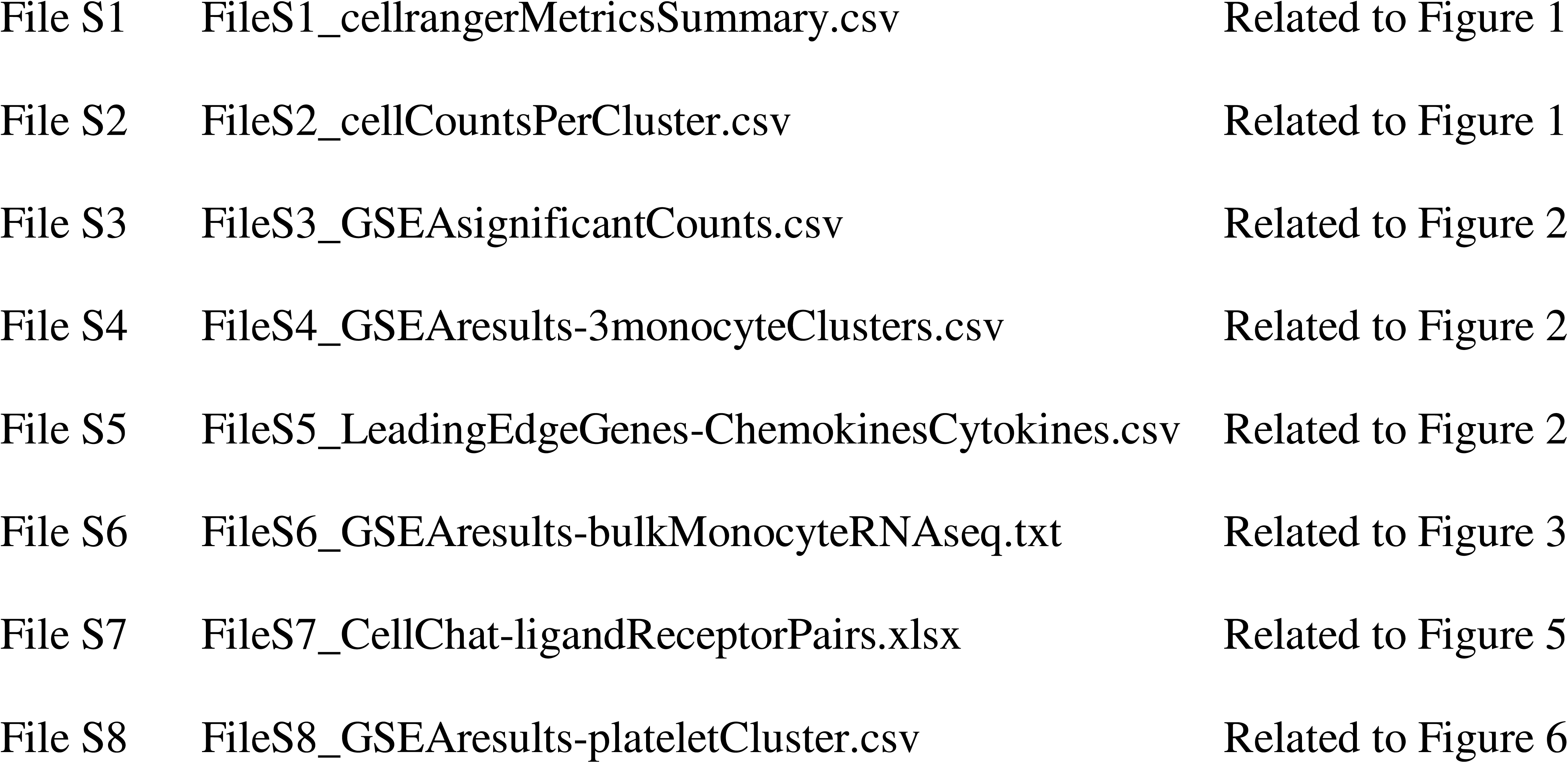

