## Supplemental figures for "Single-cell transcriptomics of the immune system in ME/CFS at baseline and following symptom provocation"

### Supplementary figure 1

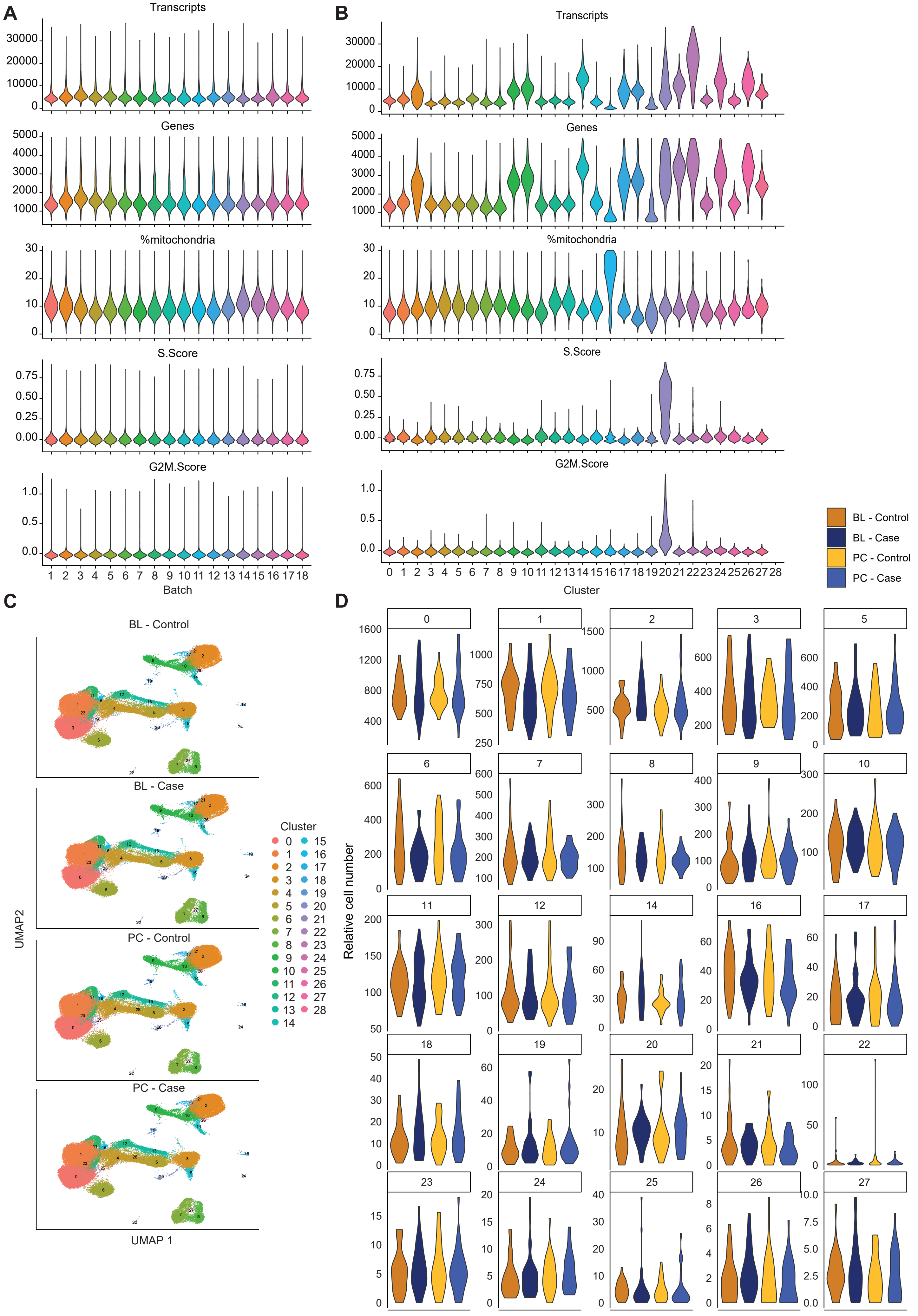

#### Supplementary figure 2

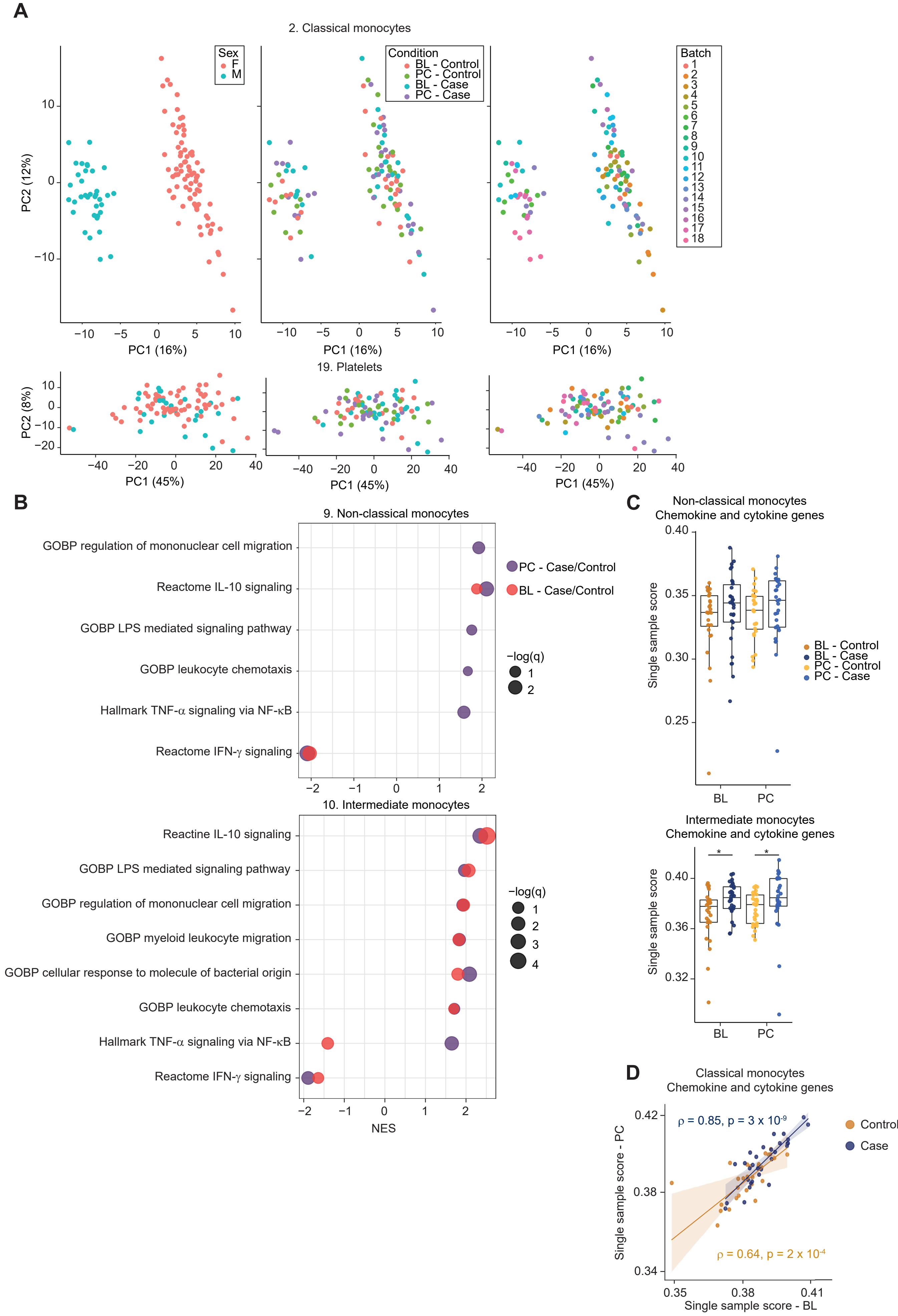

#### Supplementary figure 3

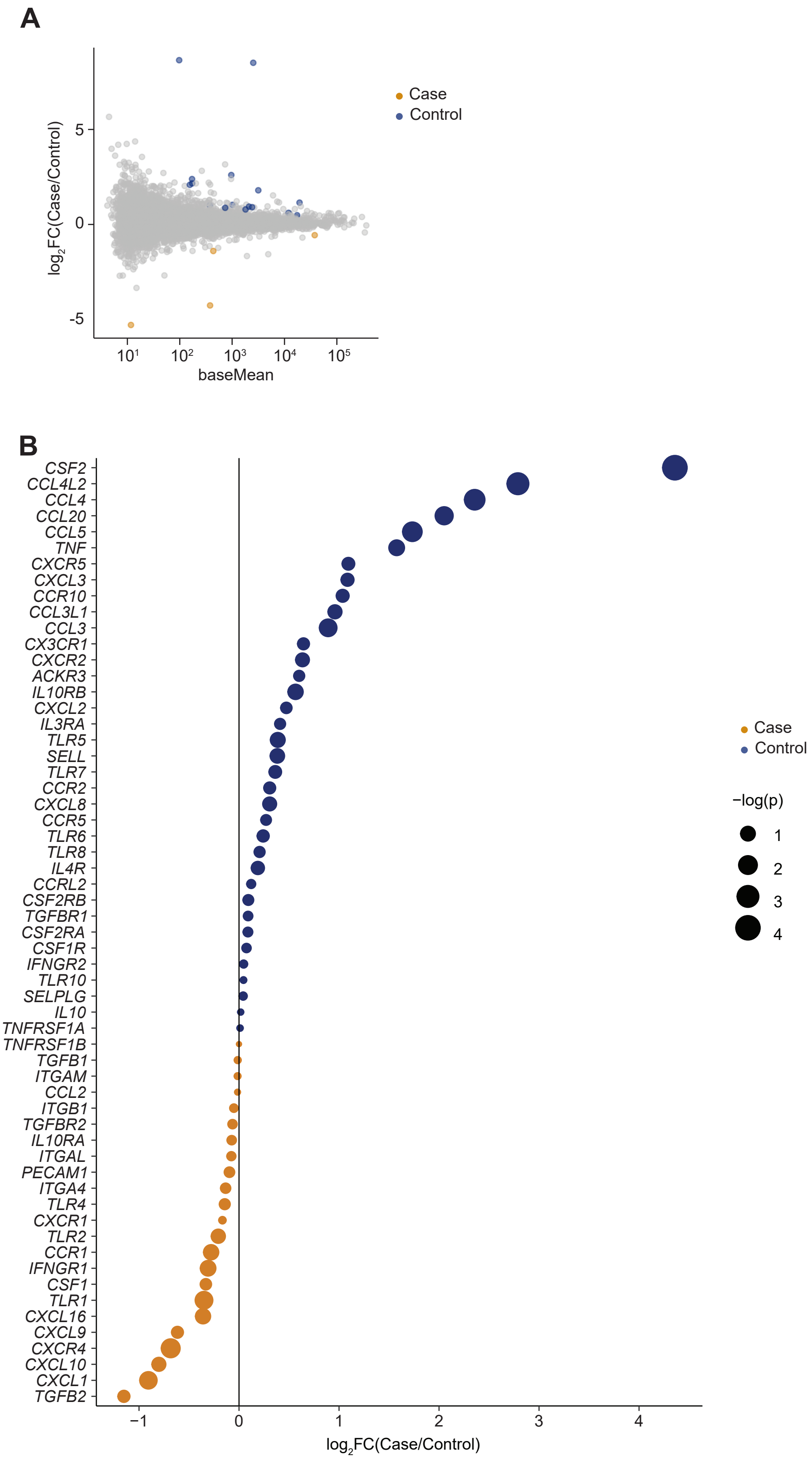

### Supplement figure 4

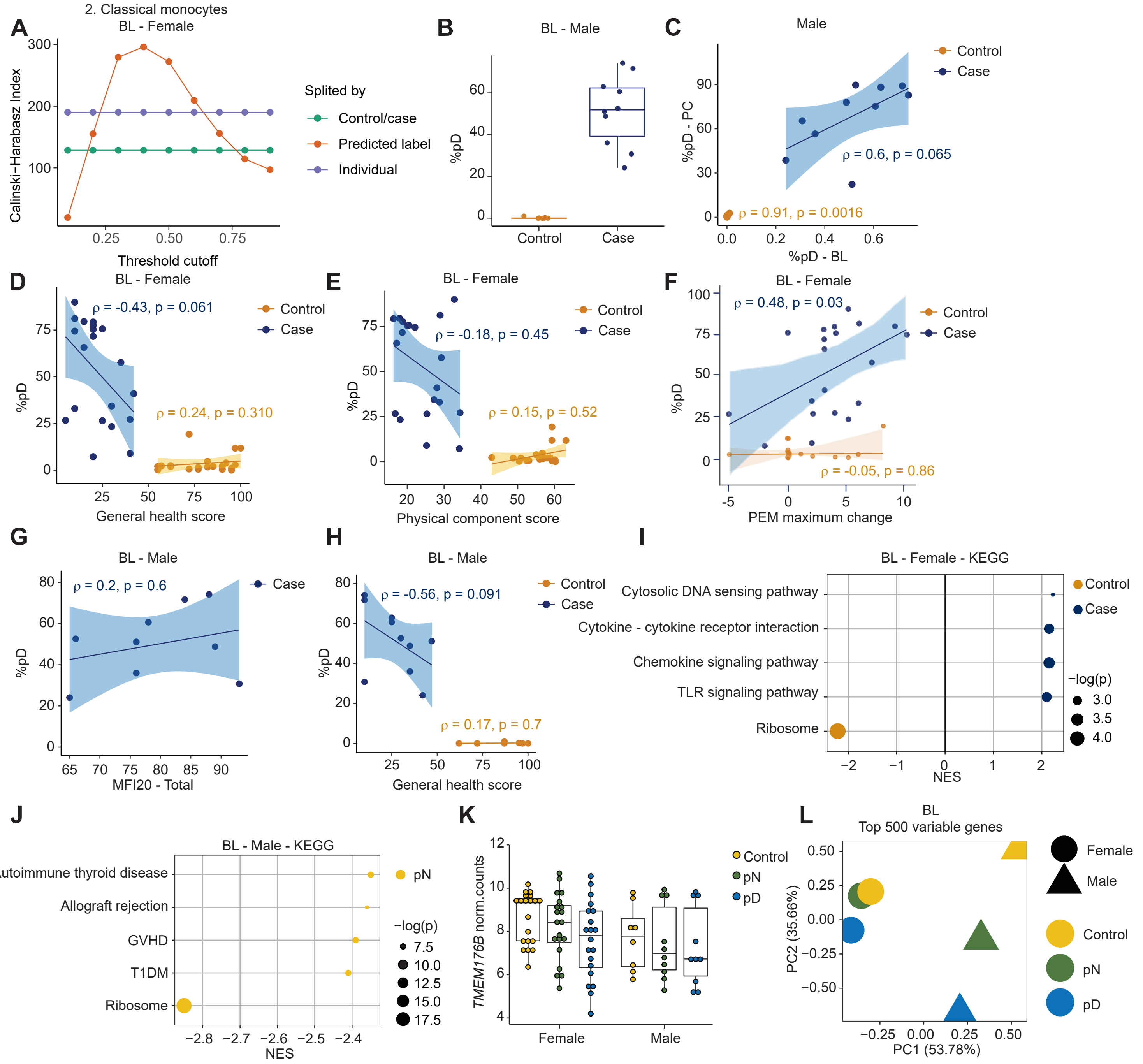

#### Supplementary figure 5

**A**

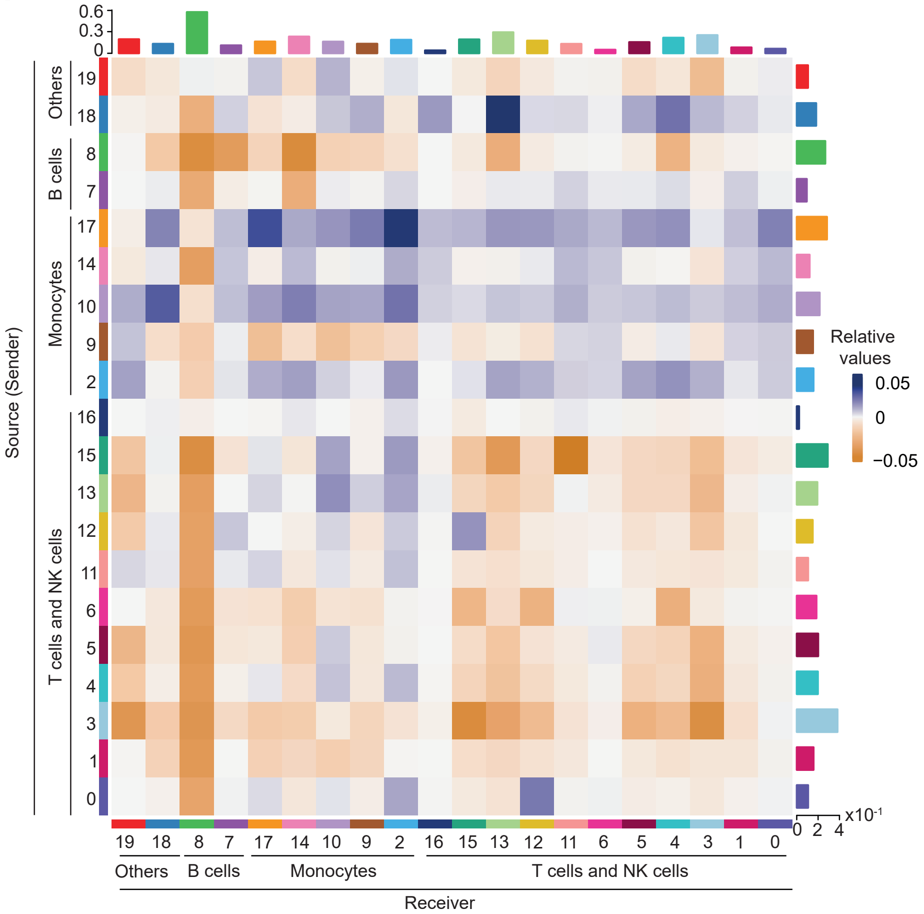


**B**

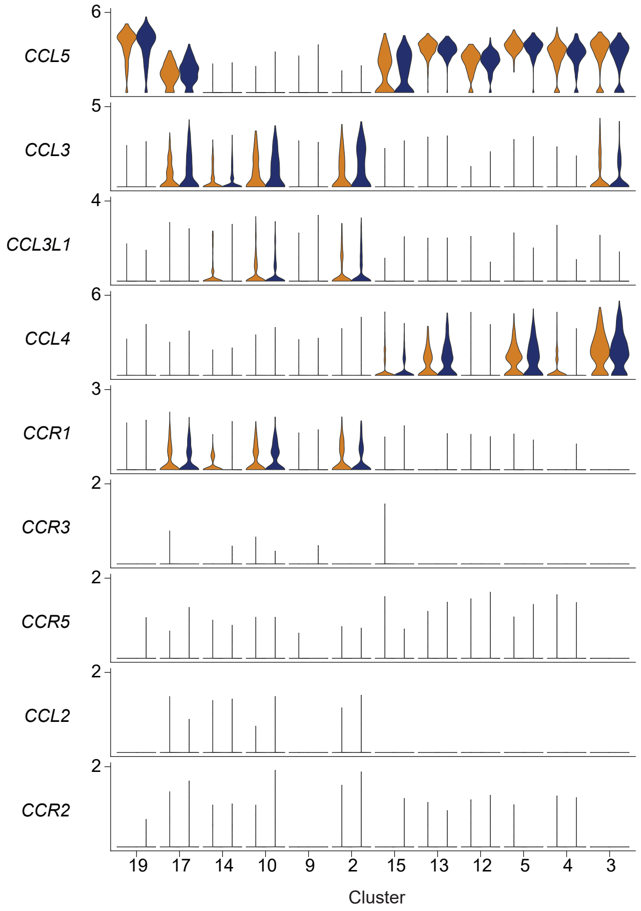
